# Modeling σ^E^ biochemical network reveals context-dependent feedback control and kinetic constraints in the envelope stress response

**DOI:** 10.1101/2025.10.22.683963

**Authors:** Cristina S. D. Palma, Natalie Allen, Martynas Basevicius, Ha Do, Kevin Li, Daniel J. Haller, Anna Konovalova, Oleg A. Igoshin

## Abstract

The bacterial cell envelope is essential for mechanical stability, barrier function, defining cell size and shape, and supporting cellular processes. Because external or internal stressors can challenge its integrity, bacteria have evolved stress response systems to maintain envelope homeostasis. For example, under stress, *E. coli* activates σ^E^ to induce a regulon that restores envelope homeostasis. The molecular mechanisms underlying the transcriptional and post-translational regulation of σ^E^ and its anti-sigma factor RseA have been well-mapped. However, how these regulatory layers function at the systems level to regulate σ^E^ activity remains unclear. Here, we combine mathematical modeling with quantitative gene expression measurements to determine how the interplay of transcription and post-translational regulation affects the dynamics of the σ^E^ response. The results suggest that the σ^E^ activity is determined by the balance between the release from RseA, resulting from its degradation, and slow binding of free σ^E^ to the intact RseA. Notably, the combined action of transcriptional and post-translational regulation reveals that autoregulation, traditionally assumed to be a positive feedback, transitions from negative to positive feedback under extreme stress, enabling a finely tuned response that prevents premature activation while ensuring a robust response under severe envelope stress. In summary, our findings elucidate the mechanisms of σ^E^ regulation, thereby advancing our understanding of how alternative sigma factor bacterial networks control stress-response pathways.

## 1. Introduction

To ensure survival in challenging environmental conditions, bacteria have evolved a variety of stress-response mechanisms that enable rapid adaptation (Bonilla, 2020; Brand et al., 2025; Gottesman, 2019; Hengge, 2014; Matin, 2009; Palma et al., 2025). Central to these responses are alternative sigma (σ) factors. Similarly to housekeeping σ-factors, alternative σ-factors can serve as subunits of RNA polymerase (RNAP) responsible for promoter recognition (Feklístov et al., 2014). However, unlike housekeeping σ-factors that keep essential gene transcription running during growth, alternative σ-factors guide RNAP to stress-responsive regulons, allowing the cell to respond and survive environmental challenges (Feklístov et al., 2014; Gruber & Gross, 2003; Helmann & Chamberlin, 1988).

The bacterial cell envelope forms the interface between the cell and its environment and is therefore the first cellular structure threatened by exposure to external stress. To preserve envelope integrity, bacteria employ a subgroup of alternative σ-factors, known as the extracytoplasmic function (ECF) family (Mascher, 2023). ECF σ-factors are used to signal extracytoplasmic conditions to the cytoplasm by activating specific regulons that maintain envelope homeostasis when cells are exposed to stresses (Brooks & Buchanan, 2008; Mascher, 2023; Raivio & Silhavy, 2001).

In *Escherichia coli*, the ECF σ-factor is σ^E^, also known as σ^24^ or RpoE. Specifically, σ^E^ coordinates the production of outer membrane proteins (OMPs) with the cell’s capacity to fold them correctly at the outer membrane. It does so by regulating the transcription of small regulatory RNAs that inhibit OMP synthesis, while simultaneously upregulating OMP folding factors in both the periplasm and the outer membrane. Additionally, σ^E^ promotes the expression of periplasmic proteases that degrade misfolded OMPs. This regulatory role is critical for preventing the toxic accumulation of misfolded OMPs, making σ^E^ essential for cell viability, even under optimal growth conditions (Konovalova et al., 2018; Yang et al., 2025). Further, σ^E^ activity is elevated in response to stressors that disrupt OMP folding (Saha et al., 2021), such as heat shock, chemical or genetic inhibition of the OMP folding pathway, and alterations in lipopolysaccharide structure. Upon activation, σ^E^ helps maintain envelope homeostasis and ensures cell survival. Notably, pre-induction or enhanced activation kinetics of σ^E^ allows cell survival under otherwise lethal conditions by allowing bacteria to adapt faster to sudden and severe outer membrane stress (Hart, O’Connell, et al., 2019; Kern et al., 2019; Konovalova et al., 2016; Leiser et al., 2012).

The post-translational regulation of σ^E^ is well-characterized (Figure 1, top panel). Under optimal conditions, σ^E^ is bound to the cytoplasmic domain of the membrane-spanning anti-σ factor, RseA (De Las Peñas et al., 1997a; Missiakas et al., 1997), resulting in low levels of free σ^E^. When the cell faces stress, σ^E^ is activated through a highly regulated proteolytic cascade (Ades et al., 1999; Alba et al., 2001; Chaba et al., 2007; Kanehara et al., 2002). First, the misfolded OMPs in the periplasm activate the membrane protease DegS, which cleaves the periplasm domain of RseA (Alba et al., 2001; Walsh et al., 2003). Next, intramembrane protease RseP cleaves the RseA transmembrane region (Kanehara et al., 2002), after which RseA is degraded in the cytoplasm, freeing σ^E^ (Flynn et al., 2004). Once free, σ^E^ can associate with the RNAP core to initiate transcription of its regulon. Of all these steps, the initial cleavage of RseA by the protease DegS has been identified as the rate-limiting step (Chaba et al., 2007). Notably, the overall degradation rate of RseA has been shown to vary across different phases of the heat-shock response, increasing approximately four-fold from optimal to stress conditions (Ades et al., 2003).

**Figure 1:**
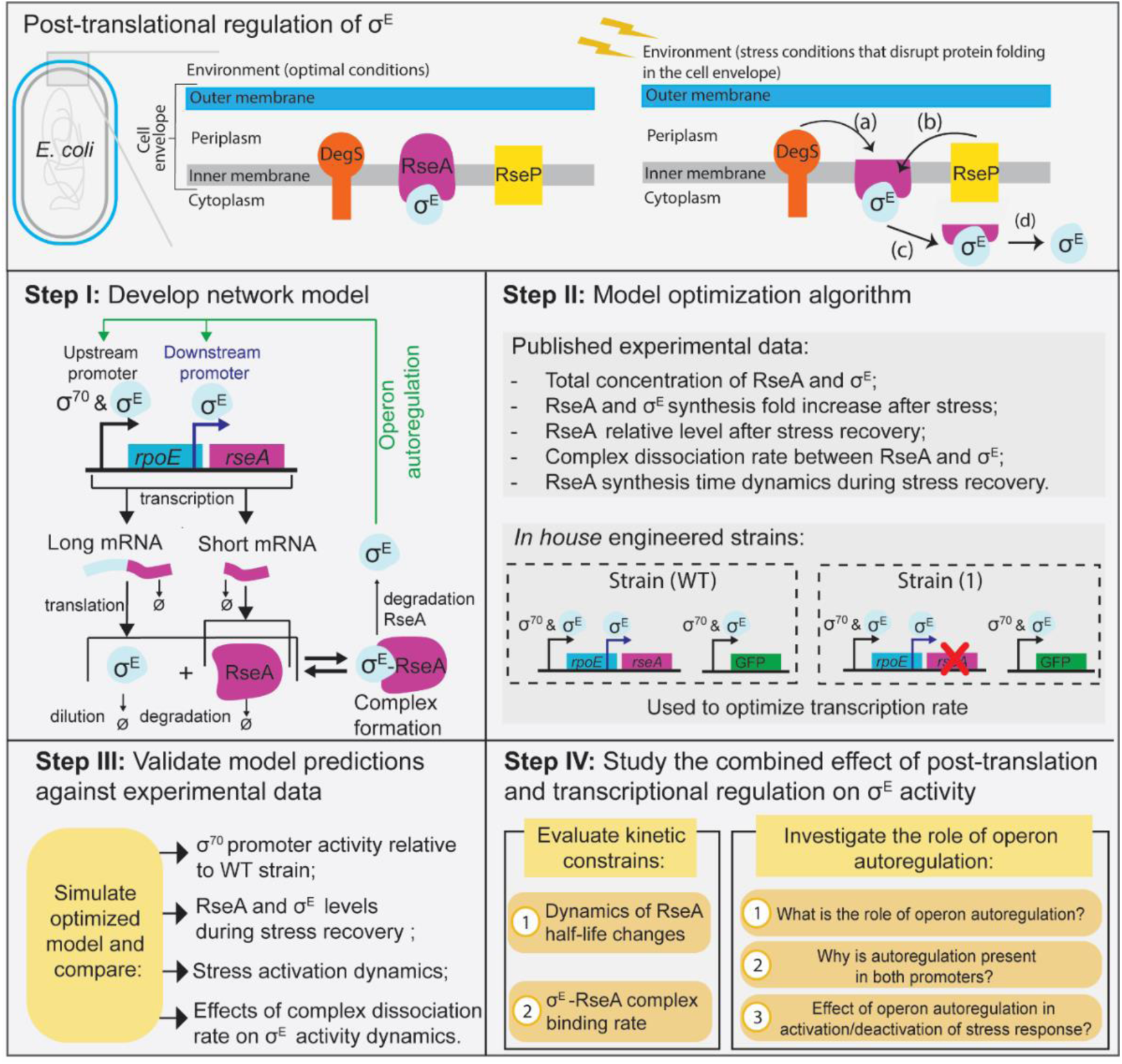
Post-translation regulation of σ^E^ and study workflow. **(Top panel)** Schematic of the *E. coli* cell envelope highlighting the key proteins involved in the post-translational regulation of σ^E^: the inner membrane–spanning anti-sigma factor RseA, and the proteases DegS and RseP. Upon envelope stress, DegS initiates the response by cleaving the periplasmic domain of RseA (a). This is followed by intramembrane cleavage of RseA by RseP (b). The resulting cytoplasmic fragment, which remains bound to σ^E^, is then released into the cytoplasm (c), where further degradation of RseA by cytoplasmic proteases frees σ^E^ (d). Adapted from (Chaba et al., 2007) **(Step I)** Illustration of the σ^E^ regulatory network. Only the key components of the *rpoE-rseABC* operon relevant to this study are represented. The upstream σ^70^ and σ^E^ specific promoter induces transcription of a long mRNA that translates to σ^E^ and RseA. A downstream σ^E^ specific promoter induces transcription of a short mRNA that translates into RseA. Post-translationally, free σ^E^ and RseA can associate into a complex. Free σ^E^ can associate with core RNAP completing the autoregulatory loop (green arrow). Notably, in σ^E^-RseA complex, RseA can suffer degradation leading to free σ^E^. **(Step II)** To estimate the unknown model parameters we implemented a model optimization algorithm. For this we made used of experimental data from (Ades et al., 2003; Collinet et al., 2000; Raina et al., 1995). In addition, we also measured promoter activity (Methods Section 4.2) of synthetically engineered strains (Methods Section 4.1) to fine tune transcriptional activity. Strain (WT) contains a reporter plasmid for the upstream promoter. In Strain (1) the gene *rseA* was knocked out from the native locus. **(Step III)** Next, we compare model predictions with experimental data (Ades et al., 2003; Chaba et al., 2007). **(Step IV)** Finally, we investigated kinetic constrains and the role of operon autoregulation in the σ^E^ regulatory network.

In addition to post-translational regulation, σ^E^ activity is also controlled at the transcription level with multiple feedback loops. Specifically, σ^E^, encoded by the *rpoE* gene, is expressed within the same operon as its anti-sigma factor RseA, the *rpoE-rseABC* operon. The expression of the *rpoE-rseABC* operon is regulated by several promoters (Klein et al., 2016; Raina et al., 1995). Two major ones include an upstream promoter recognized by both σ^70^ and σ^E^ (Missiakas et al., 1997; Raina et al., 1995), and a downstream promoter recognized only by σ^E^ (Raina et al., 1995). The upstream promoter induces transcription of a long mRNA that translates σ^E^ and RseA and other downstream genes, whereas the downstream promoter induces transcription of a shorter mRNA that only encodes RseA (Figure 1, Step I).

Despite extensive research characterizing the molecular details of the σ^E^ stress response, several key questions remain unresolved at the systems level. For example, it remains unclear how σ^E^ activity can persist when RseA is present in excess under both optimal and stress conditions (Collinet et al., 2000), and the dissociation constant between the two proteins is extremely low (less than 10 pM), as reported in (Chaba et al., 2007). Further, during stress recovery, σ^E^ activity changes substantially even though the relative levels of RseA and σ^E^ remain nearly constant (Ades et al., 2003), raising the question of which regulatory mechanism allows such behavior. In addition, the combined effect of post-translational and transcriptional regulation on σ^E^ activity (Figure 1A-green arrow) has not been investigated. Namely, it remains unclear how the combination of varying RseA degradation rates within the σ^E^ autoregulatory loop shapes σ^E^ activity across the different phases of the stress response.

To address these questions, we developed the mathematical model of the underlying biochemical network (Figure 1, Step I) following the same approaches as we did for other stress-response networks (Igoshin et al., 2007, 2006; Narula et al., 2016; Rao et al., 2021; Tiwari et al., 2010). To estimate the model parameters, we used an optimization approach (Figure 1, Step II). For this, we utilized published experimental data on total cellular levels of RseA and σ^E^ (Collinet et al., 2000), including how their relative abundances change during heat-shock recovery (Ades et al., 2003), the fold increase in their synthesis following heat-shock (Ades et al., 2003; Raina et al., 1995), and the dynamics of RseA synthesis throughout the recovery phase (Ades et al., 2003). In addition, to fine-tune the transcription rate parameters in the model, we engineered two strains and measured their relative promoter activities. Next, we validated the optimized model by comparing its predictions to corresponding experimental data (Figure 1, Step III). Finally, we use the model to predict the effect of operon autoregulation on σ^E^ activity, to identify the kinetic constraints necessary to reproduce the empirical data and to understand how the interplay of posttranslational interactions and transcriptional feedback loops shaped network dynamics (Figure 1, Step IV).

## 2. Results

### 2.1 Model construction and optimization

We began by translating the network illustration shown in (Figure 1, Step I) into a set of model reactions (Figure 2A) and mathematical equations (Methods Section 4.3, Table S1-S2). The model incorporates the details of *rpoE-rseABC* transcriptional regulation and post-translational interactions to compute concentrations of mRNA, proteins, and their complexes. Specifically, the transcription of *rseA* and *rpoE* is modeled by reactions 1 and 2 in Figure 2A. The expression of *rpoE* is regulated by an upstream (*U*) promoter. Meanwhile, the expression of *rseA* is regulated by promoter *U* and an additional downstream (*D*) promoter. As such, expression from promoter *U*, leads to the production of a polycistronic mRNA with both *rpoE* (*e*) and *rseA* (*r*) transcribed (reaction 1). Meanwhile, expression from promoter *D* produces a *rseA* mRNA (reaction 2). Translation reactions are modeled by reaction 3 and reaction 4. Post-translationally, RseA and σ^E^ interact to form a complex, denoted as *C* (reaction 5). Given that RseA degradation rate is unstable and stress-dependent (Ades et al., 2003), *C* can dissociate via RseA degradation, releasing free σ^E^. (reaction 6). Importantly, the rate of RseA degradation is the model input parameter that reflects the system stress level, as measured in (Ades et al., 2003). Finally, reaction 7 to 11 model mRNA and protein degradation/dilution.

**Figure 2:**
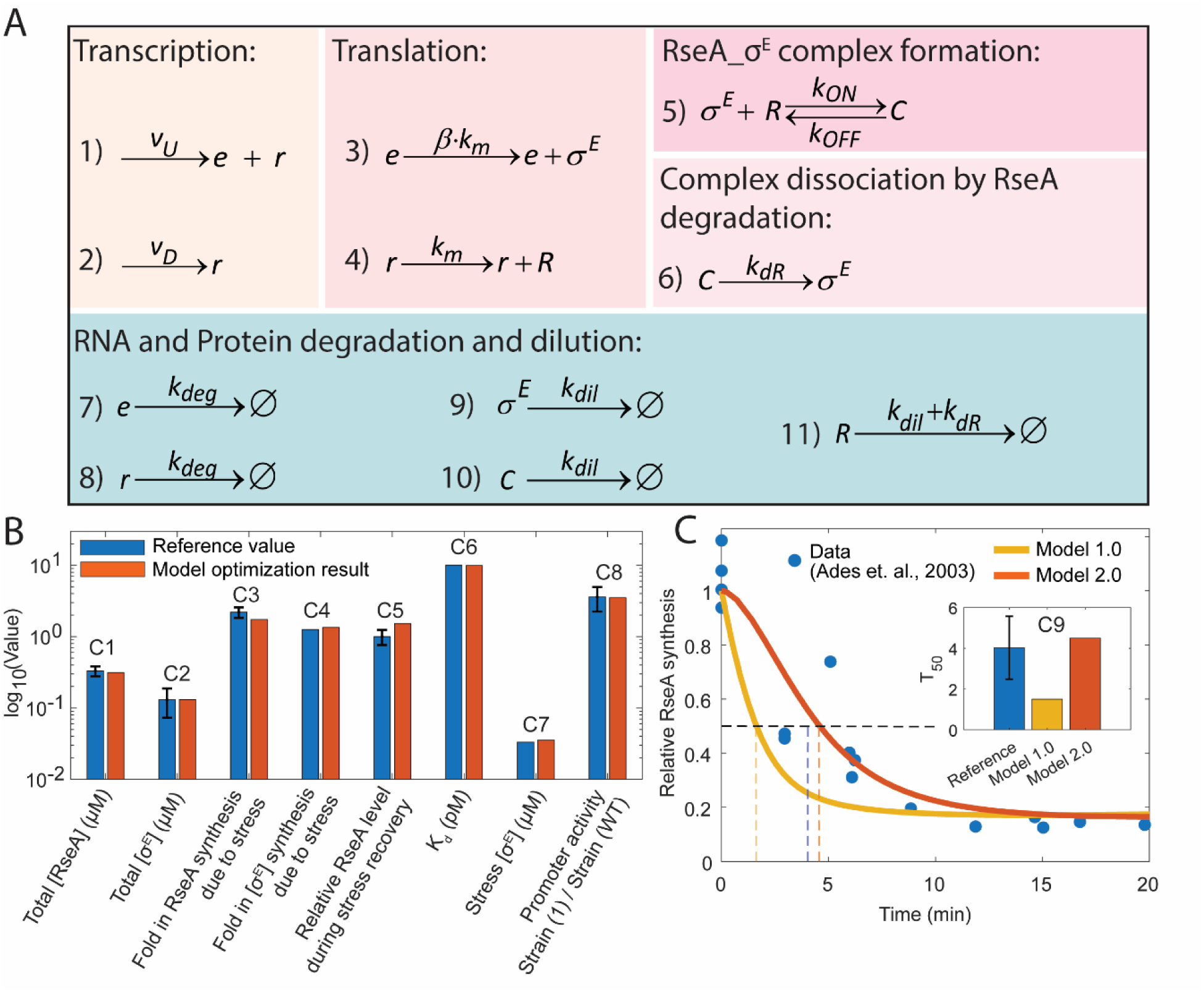
Model reactions and model optimization results. **(A)** Model reactions of the σ^E^ network. Rate constants were optimized using the optimization algorithm described in (Methods section 4.4). Transcription equations for *v_U_* and *v_D_* are defined by Eq. 1 and 2, respectively. Variables *e* and *r* represent the mRNA levels of *rpoE* and *rseA*, respectively. *R* denotes RseA protein, and *C* represents the σ^E^_RseA complex. Rate constant values are listed in Table S1. **(B)** Bar plot comparing the literature-based model constraint values (reference values) with those from the model optimization results. **(C)** Relative RseA synthesis predicted by the model, assuming an immediate change in the RseA degradation rate following a temperature downshift from 43°C to 30°C (yellow line). Blue circles are experimental data obtained from (Ades et al., 2003). Also, shown is the relative RseA synthesis predicted by *model 2.0* which assumes a gradual change in the RseA degradation rate (orange line). Horizontal black dashed line indicates 50% decay of the initial RseA synthesis value. Vertical dashed lines indicate the time it takes to reach 50% decay (T_50_) for *model 1.0* (yellow dashed), *model 2.0* (orange dashed), and reference data (blue dashed). (**Inset)** Bar plot of T_50_ values for the reference data (Ades et al., 2003), *model 1.0* (yellow line), and *model 2.0* (orange line).

Given the monomeric nature of σ-factors and the absence of cooperative binding, the transcription of *rpoE* and *rseA* is modeled using a Michaelis–Menten-like equation. Specifically, transcription from promoter *U*, which is dependent on both σ⁷⁰ and σ^E^ is described as:

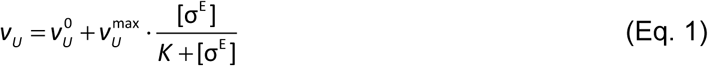

Where, 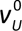 is the basal transcription rate due to σ^70^ regulation, 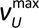 is the maximum expression rate when the promoter is saturated by σ^E^-RNAP. The variable *K* is the half-maximal binding constant. The variable [*σ ^E^*] is the concentration of the free σ^E^. Meanwhile, transcription from the downstream (*D*) promoter, a σ^E^-dependent promoter, is modeled in accordance with the following equation:

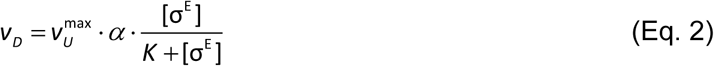

Where, 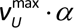 is the maximum expression rate when the promoter is saturated by σ^E^-RNAP.

To estimate the unknown model parameters, we implemented a model optimization algorithm, incorporating nine constraints to guide the fitting process (Table S3). Constrain 1 and 2 (C1 and C2 in Table S3) refer to the total concentration of RseA and σ^E^ in the cell. These values were derived from (Collinet et al., 2000), assuming a cell volume of 1fL (Phillips et al., 2013), which reports a total of 200 RseA molecules and 80 σ^E^ molecules under optimal and stress conditions. Constrain 3 (C3) refers to the ∼2.2 fold increase in RseA synthesis after subjecting cells to temperature stress (43°C, (Ades et al., 2003)). Constrain 4 (C4) refers to the ∼1.3 fold increase in σ^E^ synthesis after temperature stress (Raina et al., 1995). In addition, in (Ades et al., 2003), it is reported that during stress recovery (i.e., when conditions return to optimal), the σ^E^ activity decays but the total level of RseA remains constant. This result is considered in C5. Constrain 6 (C6), regards the reported σ^E^ and RseA dissociation constant (*K_d_*) reported in (Chaba et al., 2007). In addition, we set the concentration of free σ^E^ under stress conditions to be one-fourth of the basal total σ^E^ concentration observed under optimal conditions (C7), to prevent the model from fitting unrealistically low σ^E^ levels during stress. To further constrain the transcription rates, we engineered two strains. Specifically, we engineered a strain with a reporter plasmid for the upstream promoter (Figure 1, Step II, Strain WT, and Methods Section 4.1). The second strain also contained the same reporter plasmid but lacked the native *rseA* gene, which was knocked out from its chromosomal locus (Figure 1, Step II, Strain 1, and Methods Section 4.1). Using flow cytometer and plate reader measurements, we calculated the average relative promoter activity (Methods Section 4.2) between the two strains to be ∼ 3.6. Next, the corresponding modifications reflecting the mutant strains were implemented in the model to apply the constraint regarding their relative promoter activity (C8). Finally, to provide the model with temporal dynamics rather than relying solely on steady-state constraints, we also incorporate in C9 the reported time-course of relative RseA synthesis following stress recovery from 43 °C to 30 °C (Ades et al., 2003).

To find the best fitting rates constants that satisfy these constraints, we defined an objective function for optimization. This function minimizes the sum of the squared differences between the constraints and model predictions (Methods section 4.4). The comparison between the literature values for constraints C1–C8 and the model optimized output values is shown in Figure 2B.

Overall, constraints C1–C8 are satisfied by the optimization algorithm (Figure 2B). The poorest fit was observed for constraint C9, where the model predicted a faster decay of RseA synthesis than indicated by the experimental data (Figure 2C, yellow line versus blue circles).

### 2.2 Model of σ^E^ regulation suggests that RseA degradation rate changes gradually

In the model described in the previous section (hereafter *model 1.0*), stress recovery was assumed to cause an immediate change in the RseA degradation rate, with its half-life shifting from 4.5 minutes to 50 minutes (Ades et al., 2003). In an attempt to improve the fit of constraint C9, where the model predicted a faster decline in RseA synthesis than observed in the experimental data (Figure 2C, yellow line), we developed a revised version of the model, denoted *model 2.0*. In this version, we assume an exponential change in the RseA half-life (*T*_1/2_) from 4.5 min (*τ*_1_) to 50 min (*τ*_2_) on the scale 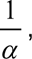 in accordance with the following equation:

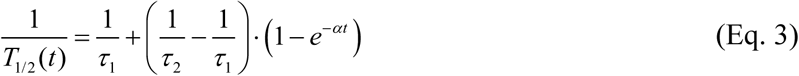

Next, we re-ran the model optimization algorithm for *model 2.0*. The results show that the model fit further improved, with the mean squared error (MSE) decreasing from 0.8 to 0.5. Notably, the fit of constraint C9 showed the greatest improvement with the RseA synthesis decay profile adjusted from a T₅₀ (i.e., the time it takes for the initial value to decay 50%) of 1.5 minutes (Figure 2C, yellow dashed line) to a slower decay with a T₅₀ of 4.5 minutes (Figure 2C, orange dashed line). This decay rate on the RseA synthesis accurately captures the experimentally observed data (Figure 2C, blue circles). We therefore conclude RseA half-life changes gradually during stress recovery, with the decay rate (*α*) in *Eq. 3* estimated by the model to be 0.005 s^-1^.

Furthermore, to assess the uncertainty of the optimized parameter values, we employed an ensemble modeling approach (Martin et al., 2023) using 2,000 independent parameter sets (Method section 4.4). Each set was locally optimized to obtain a distribution of fitted parameter values. The low variability across these distributions (Figure S1) suggests that the optimized parameter values found (Table S1) without the ensemble approach are robust.

### 2.3 Model predictions reproduce experimental data and validate proposed hypothesis

To validate the model, we compared the model predictions with experimental data on relative promoter activity, stress response activation dynamics, and total levels of RseA and σ^E^ during recovery from stress. Also, we evaluated how the complex dissociation constant (*k_OFF_*) influences the system and compared the results with the hypothesis proposed by (Chaba et al., 2007).

To quantify relative promoter activity, we engineered two other strains. Namely, *Strain (2)* carries an upstream-promoter reporter in which the σ^E^ binding site was mutated (Figure 3A, Methods Section 4.1). *Strain (3)* carries the same reporter in a native-locus *rseA* deletion background (Figure 3A, Methods Section 4.1). For each strain we measured, by flow cytometry (Methods Section 4.2), the GFP expression level relative to the wild-type (WT) strain (strain (WT) in Figure 1, Step II). As anticipated, expression level did not differ between *Strain (2)* and *Strain (3)* because the reporter in both lacks σ^E^ specificity and is therefore unaffected by the increase in free σ^E^ caused by *rseA* deletion (Figure 3B, orange and yellow bars). Absolute measurement values for each strain are shown in Figure S2B, with concordant results from plate-reader assays Figure S2C (Methods Section 4.2).

**Figure 3:**
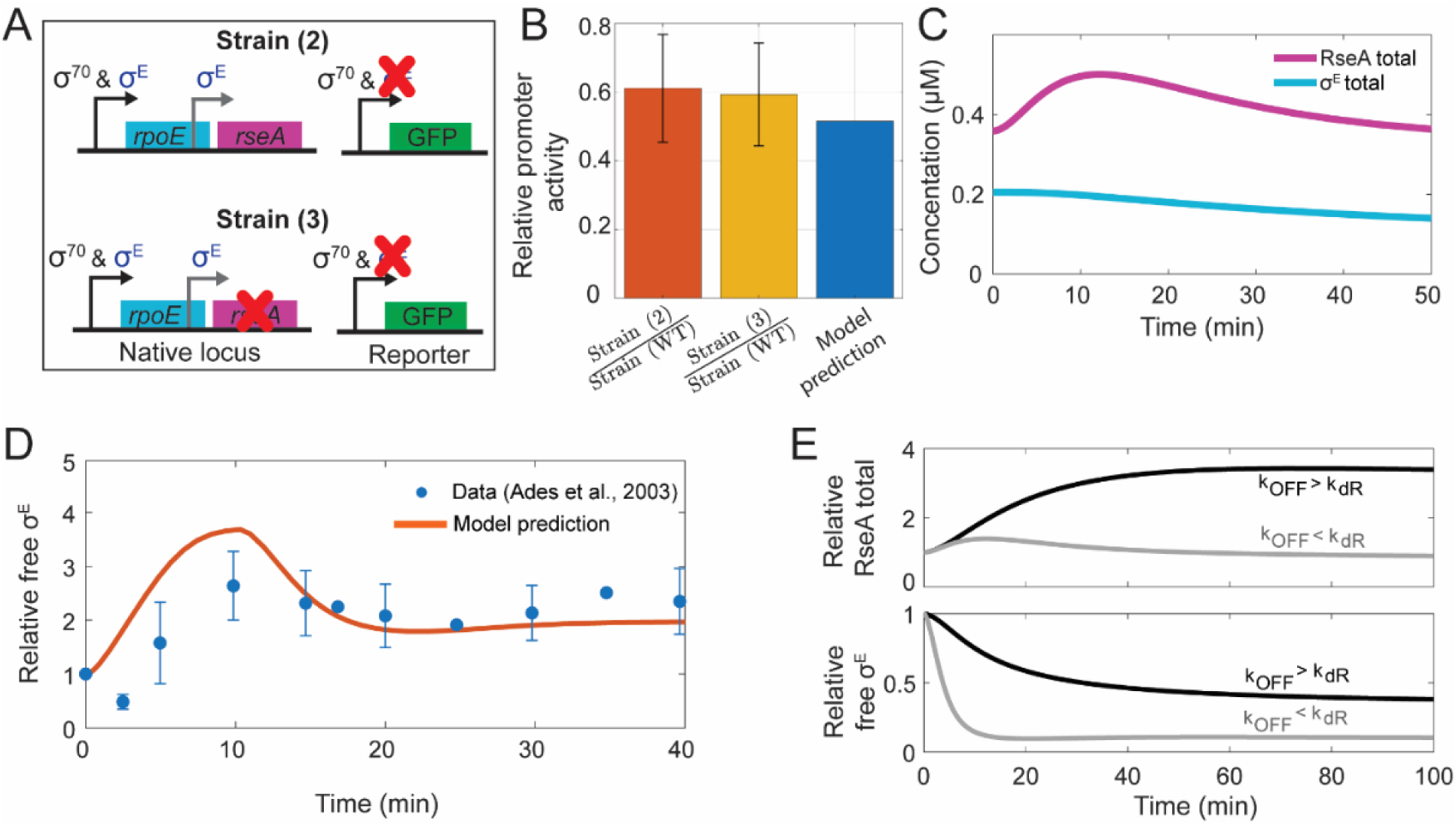
Agreement between model predictions, experimental data, and behavioral hypotheses. **(A)** Representation of the mutant strains used to calculate relative promoter activity. Strain (2) does not contain promoter specificity to σ^E^ in the reporter plasmid. In strain 3, the *rseA* was knocked out from the native locus and the reporter plasmid does not contain promoter specificity to σ^E^. **(B)** Experimental promoter activity of strain (2) and strain (3) relative to the WT strain (orange and yellow bars, respectively). Also shown is the model predicted promoter activity value (blue bar). Error bar corresponds to one standard deviation. **(C)** RseA and σ^E^ total levels predicted by the model upon stress recovery. **(D)** Model predicted stress response activation dynamics assuming a gradual decrease in RseA half-life from 8 to 2 minutes over the first 10 minutes of the stress response, followed by an adaptation phase in which the half-life stabilizes at 4.5 minutes (orange line). Also shown is the equivalent experimental data (blue dots) from (Ades et al., 2003) **(E)** Dynamics of relative RseA and free σ^E^ assuming different complex dissociation rates (*k_OFF_*). **(Upper Panel)** Relative level of RseA total during stress recovery (i.e., temperature downshift from 43°C to 30°C) assuming a complex dissociation rate (*k_OFF_*) 100-fold higher than the RseA degradation rate, *k_dR_*, (black line). Also shown is the concentration dynamics assuming a *k_OFF_* rate that is 150-fold smaller than *k_dR_*, as predicted by the model optimization algorithm (grey line). **(Lower Panel)** Relative free σ^E^ concentration during heat-shock recovery assuming *k_OFF_*rate is 100-fold higher than *k_dR_* (black line). Also shown is the concentration dynamics assuming a *k_OFF_* rate that is 150-fold smaller than *k_dR_*, as predicted by the model optimization algorithm (grey line).

We then used the model to predict the corresponding *in silico* promoter activity. Based on the model’s transcription equations (*Eq. 1* *and 2*), the relative promoter activity between these strains and the WT strain can be expressed as:

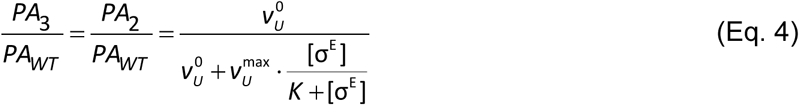

Where, *PA* stands for promoter activity. Subscripts 2/3/WT refer to strain (2), strain (3), and strain (WT), in Figure 3A and Figure 1-step II, respectively. As in *Eq. 1* and *2*, the variable 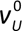 is the basal transcription rate due to σ^70^ regulation, 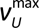 is the maximum expression rate when the promoter is saturated by σ^E^-RNAP. The variable *K* is the half-maximal binding constant. The variable [*σ ^E^*] is the concentration of the free σ^E^. The results demonstrate that the model accurately predicts the relative promoter activity (Figure 3B, blue bar).

In addition, the model also predicts that upon stress recovery, the total RseA and σ^E^ levels remain relatively constant over time (Figure 3C). This prediction aligns well with experimental observations reporting that RseA and σ^E^ levels vary by no more than ∼1.5-fold, while σ^E^ activity drops rapidly by ∼8-fold during the recovery phase. Furthermore, we found that the relatively constant levels of RseA and σ^E^, despite the decline in σ^E^ activity, is explained by the coordinated decrease in both their production and degradation rates, which preserves their steady-state levels (Figure S3).

Next, we compared the model-predicted stress response activation kinetics with the experimental data of RseA synthesis dynamics reported in (Ades et al., 2003). Notably, a step change in the RseA degradation rate during the activation of the stress response does not accurately capture the experimental data (Figure S4). However, when we instead assume a gradual change in RseA half-life, using the same decay rate estimated by the model optimization algorithm (Results Section 2.1), the simulated activation dynamics closely matches the experimental observations (Figure 3D, orange line). Overall, we conclude that the RseA half-life changes gradually not only during stress response deactivation (Results Section 2.2) but also during activation, following the same decay rate (Eq. 3). In addition, the model-predicted σ^E^ activity, across all phases of the stress response, shows a strong agreement with the experimental data reported by (Ades et al., 2003) (Figure S5).

Finally, we compared how the complex dissociation rate constant (*k_OFF_*), relative to the RseA degradation rate (*k_dR_*) determines the system’s behavior, and compared the results with the hypothesis proposed in (Chaba et al., 2007) that during stress recovery the rapid change in σ^E^ levels can be explained if *k_OFF_* is much smaller than *k_dR_*. Otherwise, the σ^E^ levels would not decrease rapidly but would instead decline in concert with the increase in RseA.

To determine whether our model can reproduce this hypothesis, we performed deterministic simulations for each of the two cases described above: 1) *k_OFF_* higher than *k_dR_*, and 2) *k_OFF_* lower than *k_dR_*. In accordance with (Chaba et al., 2007), the model predicts that a fast dissociation rate does not result in rapid decay of free σ^E^ levels, instead it causes a decrease in concert with the increase in RseA total levels (Figure 3E, black lines). Conversely, a slow dissociation rate enables a fast decrease in free σ^E^, while maintaining relatively constant total RseA levels (Figure 3E, grey lines).

To understand the mechanism behind this behavior, we evaluated the dynamics of free σ^E^ production for the two cases. In case 1, we found that the production of free σ^E^ is primarily controlled by the rate constant k_OFF_, as such changes in the RseA degradation rate do not affect the production rate of free σ^E^. Nevertheless, the total levels of RseA will increase gradually to reach a new steady state due to its slower degradation. Consequently, the free σ^E^ degradation term *k_ON_·[R]* (E3 in Table S2) increases in concert leading to a decrease in free σ^E^ that follows the dynamics of the RseA total levels increase. Meanwhile, if *k_OFF_* is small the amount of free σ^E^ in the system is controlled by the RseA degradation rate. As such, if the RseA degradation rate increases rapidly, as in stress recovery, changing from a 4.5 to a 50-minute half-life (Ades et al., 2003), the production of σ^E^ will be rapidly decreased leading to the fast σ^E^ decay observed.

### 2.4 The slow rate of σE–RseA complex formation keeps a large fraction of σE free

Studies have shown that the *in vitro* affinity between σ^E^ and the cytoplasmic domain of RseA is very high, with an equilibrium dissociation constant (K_d_) smaller than 10 pM (Chaba et al., 2007). It has been also reported that the cellular ratio of RseA to σ^E^ is about 5:2 and remains unchanged, even when σ^E^ activity increases in response to heat-shock stress (43 °C) (Collinet et al., 2000). This raises the question of how, under these conditions, free σ^E^ can exist in the cell at levels sufficient not only to maintain basal activity, which is essential for growth, but also to further activate its regulon during stress.

To address this question, we examined the model optimized parameters. The optimized model suggests that the rate of σ^E^–RseA complex formation (*k_ON_*) is lower than previously estimated from *in vitro* experiments, with a value of 0.08 µM⁻¹s⁻¹ compared to the reported 15 µM⁻¹s⁻¹ (Chaba et al., 2007). To assess whether excess RseA can only occur for small *k_ON_* values, we conducted two tests. First, using the optimized model, we evaluated the relative amount between total RseA and total σ^E^ for different *k_ON_* values. The results show that excess of RseA is only possible for *k_ON_* < 1 µM^-1^s^-1^ and an excess of 2.5-fold requires a *k_ON_* < 0.1 µM^-1^s^-1^ (Figure 4, orange line). In the second test, we tested different *k_ON_*values, but for each tested value, we allow for parameter re-optimization to see if any other combination of parameters allows an excess of RseA. The results suggest that even with re-optimization, an excess of RseA is not possible for *k_ON_* values > ∼1 µM^-1^s^-1^ (Figure 4, blue line). Altogether, these results suggest that, free σ^E^ can exist in the cell even under high σ^E^ -RseA affinity and excess of RseA (Chaba et al., 2007; Collinet et al., 2000), provided that *k_ON_* is lower than the value estimated from *in vitro* measurements.

**Figure 4:**
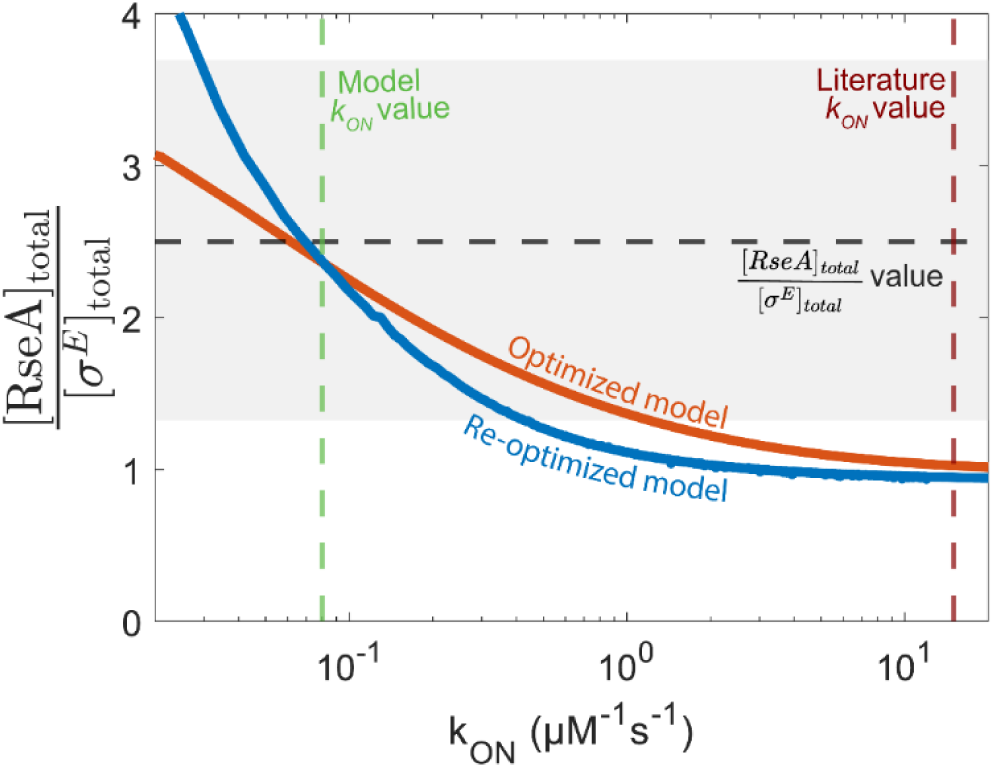
Ratio between the total levels of RseA and σ^E^ for different complex formation rates (*k_ON_*), assuming *k_OFF_* / *k_ON_* = 10 pM (Chaba et al., 2007). All other model parameters were kept fixed (orange line). Also shown are the results when model is re-optimized for each *k_ON_* value (blue line). The dashed black line indicates the literature-reported fold excess of RseA relative to σ^E^ (Collinet et al., 2000). The dashed green line indicates the model optimized *k_ON_* for which MSE is minimized. The red dashed line indicates the literature value of *k_ON_*(Chaba et al., 2007). Shadow area corresponds to the standard deviation of the excess of RseA relative to σ^E^.

### 2.5 σ^E^ feedback gain is negative during optimal growth conditions and becomes positive when cells face extreme stress

One important question that this model can help investigate is the effect of operon autoregulation and the effects of the increased *rpoE-rseABC* transcription on the σ^E^ activity.

For example, our previous work on networks integrating transcriptional and post-translational regulation has shown that positive autoregulation of sigma factors and two-component system response regulators can result in overall negative feedback when it also drives the expression of negative posttranslational regulators (Narula et al., 2016; Rao & Igoshin, 2021; Ray & Igoshin, 2010; Ray et al., 2011).

To determine whether this phenomenon also occurs in the σ^E^ regulatory network, we constructed an ‘open-loop’ version of the model in which the autoregulatory feedback of σ^E^ was removed (Figure 1A, Step I, green arrow). In this configuration, operon expression is no longer regulated by σ^E^ but instead by an external input, S₀, which is set to be a tunable model parameter. Mathematically, this is implemented by replacing the concentration of free σ^E^ in the transcriptional functions (*Eq. 1* *and 2*) with *S_0_.* Next, to assess the feedback strength, we first simulate the full (closed-loop) model and record the steady-state concentration of free σ^E^. We then assign this value to S₀ in the open-loop model to ensure that all molecular species and complexes are initialized at the same steady-state concentrations in both models. Finally, we perturb S₀ by 5% and compute the resulting steady states to estimate the loop gain of the feedback system.

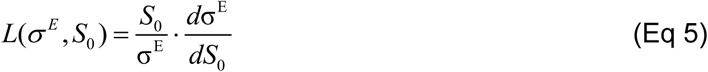

The sign of *L* defines the sign of the overall feedback loop, *i.e.* whether increases in the operon transcription will increase (+) or decrease (-) the σ^E^ transcriptional activity. The absolute value characterizes the network sensitivity to changes in transcription. Our results show that feedback is negative at optimal growth conditions and becomes positive for extreme stress conditions (Figure 5A, bottom panel).

**Figure 5:**
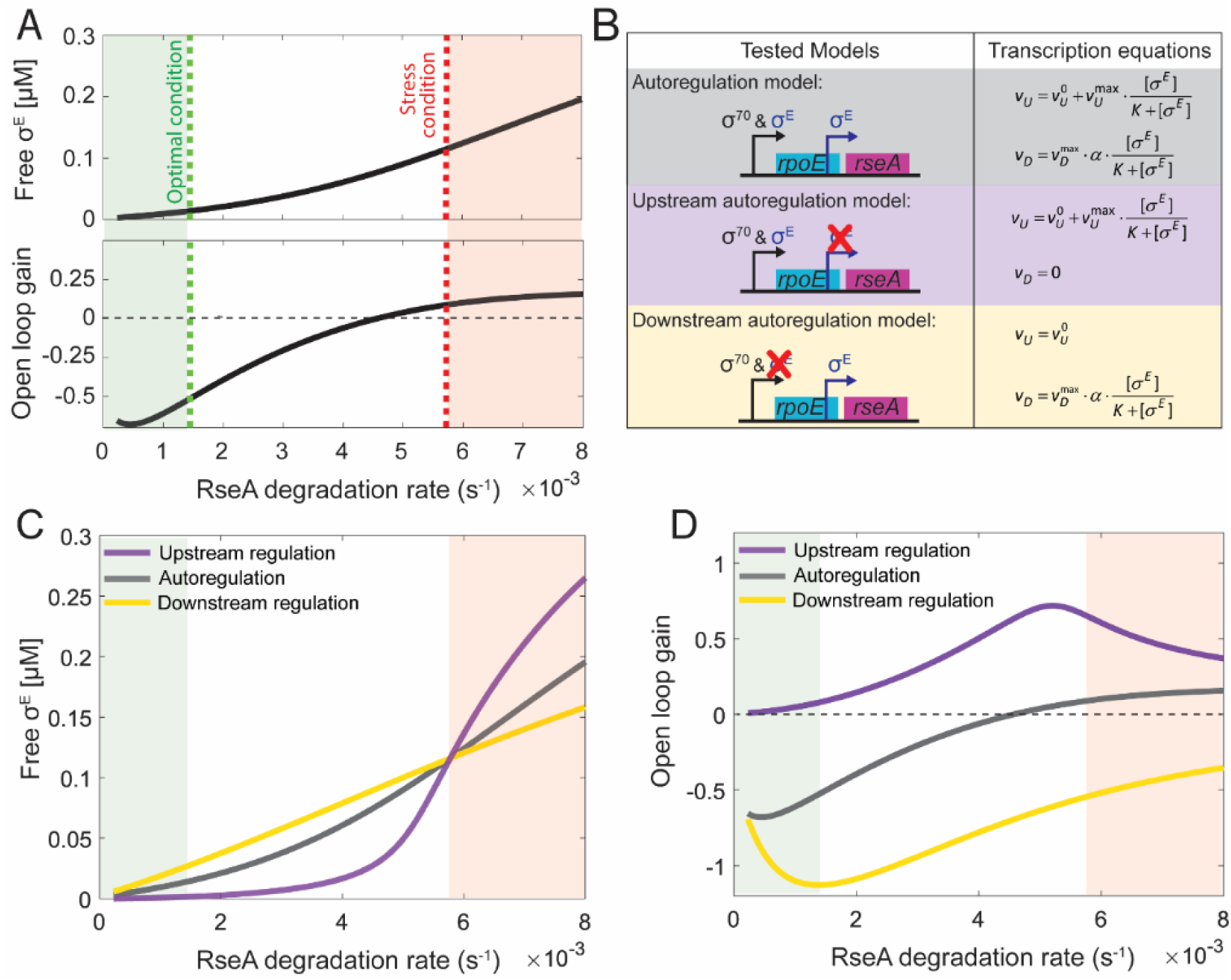
Analysis of operon autoregulation. **(A) Top panel:** Free σ^E^ concentration as a function of RseA degradation rate. **Bottom panel:** Open loop gain as a function of RseA degradation rate. Open loop gain was calculated according to *Eq. 5*. The green shaded area indicates the range of RseA half-life values under non-stress conditions. The vertical green dashed line marks the upper boundary of this range, corresponding to a half-life of 8 min (Ades et al., 2003). The red shaded area indicates the range of RseA half-life values under stress conditions. The vertical red dashed line marks the lower boundary of this range, corresponding to a half-life of 2 minutes (Ades et al., 2003). **(B)** Summary illustration of the tested models and the corresponding modifications made to the model transcription equations. Three models were tested. The ‘autoregulation model’ refers to the complete model where operon autoregulation is kept unaltered. In the ‘upstream autoregulation model’, the downstream σ^E^-specific promoter was deleted, making σ^E^ feedback to act only through the upstream promoter. In the ‘downstream autoregulation model’, the upstream σ^E^ regulation was deleted, making σ^E^ feedback to act only through the downstream promoter. Regulation by σ^70^ on the upstream promoter was kept unaltered. Also shown are the transcription reactions for *rpoE* (e) and *rseA* (r), along with the corresponding modifications made to the transcription equations for each model (Methods Section 4.3). **(C)** Free σ^E^ concentration as a function of RseA degradation rate for each of the models described in panel B. **(D)** Open loop gain as a function of RseA degradation rate for each of the models described in panel B. Open loop gain was calculated according to *Eq.* 5.

### 2.6 Switch between negative to positive feedback gain requires σ^E^ feedback in both promoters

To investigate the specific contribution of operon autoregulation at each promoter, we developed models in which σ^E^ feedback was included at only one promoter at a time (Figure 5B). In the first model (hereafter, the upstream autoregulation model) both *rpoE* and *rseA* are regulated solely by the upstream σ^70^ and σ^E^ dependent promoter. In the second model (hereafter, the downstream autoregulation model) *rpoE* is regulated solely by a σ^70^ promoter, while *rseA* is regulated by the σ^E^ promoter. The corresponding modifications to the model’s transcription reaction equations for each case are also shown in Figure 5B.

The results show that the steady-state levels of free σ^E^ are not significantly affected by the presence of σ^E^ mediated feedback in both promoters, or its presence solely at the downstream promoter (Figure 5C, grey and yellow lines, respectively). Meanwhile, the presence of σ^E^ feedback solely at the upstream promoter results in lower steady-state levels at low RseA degradation rates. However, as the degradation rate increases, the steady-state levels surpass those of the model with feedback in both promoters (Figure 5C, purple versus grey line). This suggests that restricting σ^E^ feedback to the upstream promoter leads to an amplified response under high RseA degradation conditions.

Next, we evaluated the effects of operon autoregulation at each promoter on the feedback gain. The results show that when feedback is present only at the upstream promoter, the gain is in the positive regime. In contrast, when feedback is restricted to the downstream promoter, the gain shifts to the negative regime (Figure 5D). These findings indicate that autoregulation must be present at both promoters to enable a switch between negative and positive feedback regimes.

### 2.7 σ^E^ autoregulation slows down the stress-response activation dynamics

To understand how feedback loops affect the dynamics and the steady state free σ^E^ stress response levels, we compared the results of the model with those of the system with no autoregulation. To this end, we considered a scenario in which transcription of the *rpoE-rseABC* operon is constitutive, with the rate set to the steady-state production level observed under stress conditions, corresponding to an RseA half-life of 2 minutes (Ades et al., 2003). We then varied the RseA degradation rate while keeping transcription constant and examined how these changes impact the steady-state concentration of free σ^E^.

Our results show that removing the feedback does not significantly alter the steady-state concentration of free σ^E^ (Figure 6A). Next, we investigated how the feedback existence affects the stress response activation and deactivation dynamics. To test activation dynamics, we changed RseA half-life from 8 min to 2 min at t=0 h for both the complete model (i.e. with feedback present) and the model for which transcription is constitutive (Figure 6B). To test for deactivation dynamics, we changed RseA half-life from 2 min to 8 min at t=0 h (Figure 6C). The results show that feedback slows activation dynamics (Figure 6B), requiring ∼250 min for the system with feedback to reach the steady-state free σ^E^ levels observed without feedback (Figure 6B, inset). Notably, the deactivation dynamics are unaffected by the presence of feedback (Figure 6C), with the steady state of each condition reached within ∼20 min. However, as seen in Figure 6A, the steady state without feedback is slightly lower than that with feedback. Together, these findings suggest that feedback may serve as a safeguard, ensuring that the stress response is engaged only mildly when triggered by transient environmental stresses, while still allowing a rapid deactivation once the perturbation is resolved.

**Figure 6:**
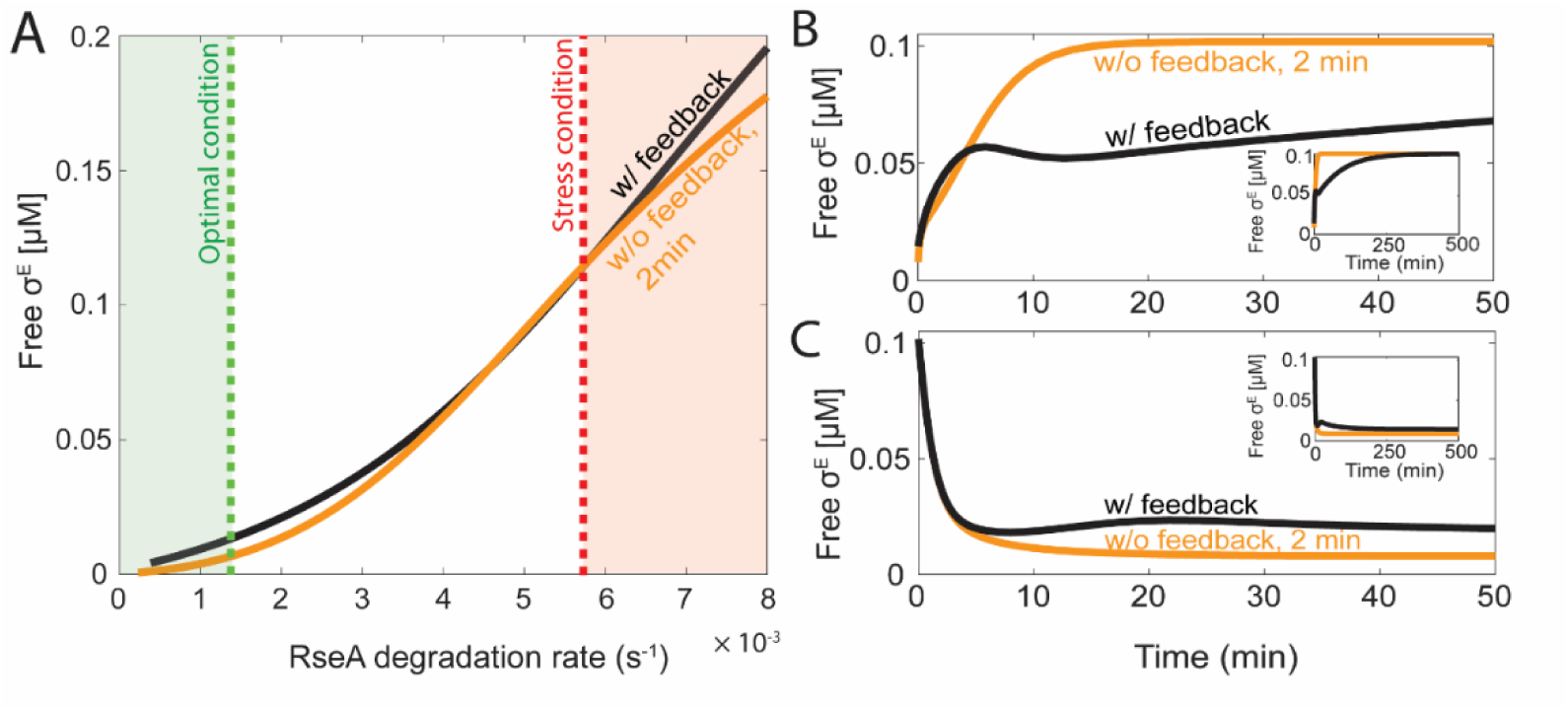
Effects of σ^E^ autoregulation in stress response activation and deactivation dynamics. **(A)** Free σ^E^ concentration in the presence (black line) and absence of operon autoregulation (yellow line). In the absence of operon autoregulation, the transcription rate was set to be constitutive and equal to the steady state transcription rate corresponding to an RseA half-life of 2 min. **(B)** Stress response activation dynamics in the presence (black line) and absence (yellow line) of operon autoregulation. For the absence of operon autoregulation, the transcription rate was set to be constitutive and equal to the steady state transcription rate corresponding to an RseA half-life of 2 min. In both scenarios, stress was simulated by shifting the RseA half-life from 8 min to 2 min at t = 0 h. The inset shows a zoomed-out view with the maximum time extended to 500 min.**(C)** Stress response deactivation dynamics under the same conditions as in graph C. To simulate the end of stress, the RseA half-life was shifted from 2 to 8 minutes at t = 0 h. The inset shows a zoomed-out view with the maximum time extended to 500 min.

## 3. Discussion

During optimal growth and stress conditions, the alternative sigma factor σ^E^ prevents accumulation of toxic misfolded outer membrane proteins (Konovalova et al., 2018). The activity of σ^E^ is regulated post-translationally through proteolytic degradation of its anti-σ factor RseA (Ades et al., 1999). Further, the expression of σ^E^ and RseA is also regulated at the transcriptional level. To our knowledge, we have developed the first mathematical model of the σ^E^ regulatory network that integrates both transcriptional and post-translational layers of regulation. The model accurately predicts the promoter activity of mutant strains (Figure 3B), the relative total levels of RseA and σ^E^ upon stress recovery (Figure 3C), the σ^E^ stress activation kinetics (Figure 3D), and validates the literature predictions of how the complex dissociation constant influences the system (Figure 3E).

Beyond reproducing published results, the model also provides novel insights. While it has been reported that RseA degradation changes upon stress (Ades et al., 2003), the temporal dynamics of this change remained unknown. Our model suggests that both upon stress and during the recovery phase, the RseA degradation rate evolves gradually (Figure 2C), following an exponential trend with a rate constant of approximately 0.005 s⁻¹. Given that the RseA degradation rate controls the timing of σ^E^ activation, degrading RseA too quickly could cause σ^E^ to become overactivated (Figure S4), leading to an excessive and potentially detrimental stress response. On the other hand, if degradation is too slow, the cell may fail to respond effectively to stress. Thus, a gradual change in RseA degradation enables an adaptive controlled response ensuring timely σ^E^ activation/deactivation.

While autoregulation is often interpreted as a positive feedback mechanism, our results challenge this assumption by revealing that its functional role is context-dependent. Specifically, under slow RseA degradation rates, typical of optimal conditions, autoregulation behaves as a negative feedback loop, decreasing σ^E^ activation. Conversely, under high RseA degradation rates, typical of stress conditions, the same regulatory architecture amplifies σ^E^ activation, functioning as a positive feedback loop (Figure 5A). This is explained by the fact that under optimal conditions, autoregulation increases both σ^E^ and RseA, but because RseA is long-lived and binds σ^E^ with high affinity, most of the additional σ^E^ becomes sequestered. As a result, the net effect is a reduction in free σ^E^ levels, which makes autoregulation behave as negative feedback. In contrast, under stress conditions, RseA is degraded rapidly, so the RseA produced is quickly eliminated while σ^E^ accumulates, leading to amplification of σ^E^ activity as positive feedback. This regulatory mechanism allows cells to buffer σ^E^ activity against minor fluctuations in σ^E^ levels under optimal conditions, for example due to stochastic bursts of transcription. In contrast, under stress conditions, it allows for an amplifying role, enhancing σ^E^ activity to ensure the activation of the stress response.

Moreover, we also found that the existence of σ^E^ autoregulatory loops help prevent overactivation of the envelope stress response (Figure 6B). As such, one can conclude that a key physiological role of the σ^E^ autoregulatory loops is to prevent inappropriate activation of the stress response, particularly under fluctuating environmental conditions where transient signals might otherwise trigger unnecessary activation, which can be toxic (Mitchell & Silhavy, 2019).

Another unresolved question in the literature regarding the σ^E^ network is the observation that σ^E^ exhibits basal activity and is essential under optimal conditions (De Las Peñas et al., 1997b). However, it has also been reported that σ^E^ and RseA exhibit strong binding affinity (Chaba et al., 2007), and that RseA is present in excess relative to σ^E^ under non-stress and stress conditions (Collinet et al., 2000). As such, one would expect that under these conditions, all σ^E^ would be sequestered through complex formation with RseA, preventing at least basal σ^E^ activity. However, our model suggests that complete sequestration does not occur because the σ^E^–RseA complex formation rate is slower than previously estimated from *in vitro* measurements. This discrepancy could be explained by the fact that the *in vitro* measurements were performed using the cytoplasmic domain of RseA. Overall, this finding reveals a previously unrecognized regulatory regime in σ-factor networks. Traditionally, these networks are understood to operate through a stoichiometric ratio between the σ-factor and its anti-σ factor, where an excess of anti-σ factor suppresses σ activity (Narula et al., 2016). Our results indicate that σ^E^ operates differently, with its activity being governed not by fixed stoichiometric ratios but by kinetic control, with slow binding to RseA and slow release through RseA degradation determining the timing and extent of activation.

In the model developed here, the influence of σ^E^ on the rate of RseA degradation is implemented phenomenologically, meaning that we do not represent the underlying mechanistic cause, but instead reproduce its observable effect on the RseA degradation rate. This choice is motivated by the fact that the observable behavior has been explicitly measured (Ades et al., 2003), whereas the mechanistic details, although known to depend on the amount of unfolded outer membrane proteins, remain unclear. In the future, it would be valuable to expand the model to directly integrate the σ^E^ feedback on RseA degradation, enabling evaluation of how changes in this pathway influence the overall stress response during stress adaptation and how it is tuned by different types of stress.

Our model demonstrates that transcriptional control of the *rpoE-rseABC* operon can significantly alter the behavior of the σ^E^ regulatory network. In this work, we focused on the regulatory influence of the two major operon promoters. However, several minor upstream promoters of *rpoE-rseABC* are regulated by diverse factors, including the general stress response alternative sigma factor RpoS, multiple envelope-associated two-component systems and stress pathways (Klein et al., 2016). While the functional consequences of this transcriptional regulation on σ^E^ activity remain unclear, it likely represents a key integration point for distinct stress responses within a higher-order regulatory network that coordinates envelope homeostasis.

Overall, our study advances the understanding of how the bacterial cell envelope, the most complex and essential cellular structure (Saha et al., 2021), is maintained under stress conditions. This is particularly important given that the envelope biogenesis and outer membrane protein assembly pathways are major targets for antibiotic development (Bisht et al., 2024; Storek et al., 2024). σ^E^ envelope stress response allows bacteria to survive otherwise lethal perturbations in the outer membrane (Hart, O’Connell, et al., 2019; Kern et al., 2019; Konovalova et al., 2016; Leiser et al., 2012). σ^E^ pathway is expected to emerge as a resistance mechanism to outer membrane biogenesis inhibitors, and is itself a target for developing antimicrobials that prevent bacterial survival by blocking key adaptation steps, thereby serving as an anti-resistance strategy (Bongard et al., 2019; Hart, Mitchell, et al., 2019; Konovalova et al., 2018). Thus, elucidating the regulatory mechanisms behind σ^E^ and other stress responses is necessary to provide critical insight into how bacteria maintain envelope integrity and promote cell survival.

## 4. Methods

### 4.1 Bacterial strains and growth conditions

All the bacterial strains used in this study are listed in Table S4 and derived from MC4100. Strains were grown at 37° in Lysogeny broth (LB) [10 g/L tryptone, 5 g/L yeast extract, 10 g/L NaCl], supplemented when appropriate at the following concentrations of antibiotics: chloramphenicol 20 μg/mL, kanamycin 25 μg/mL.

To construct transcriptional reporter fusions, gene blocks corresponding to the native promoter region of the *rpoE-rseABC* operon and its mutant derivative with a mutated σ^E^ binding site (synthesized by Eurofins Genomics) were cloned upstream of the *gfpmut3* gene using NEBuilder (NEB) into the low-copy-number vector with p15A ori and a chloramphenicol resistance cassette. The complete sequences of P*rpoE* (σ70 & σE) and P*rpoE* (σ70 only) are in Table S5.

### 4.2 GFP fluorescence measurement and promoter activity calculation

For flow cytometry measurements (Figure 3B and Figure S2B), overnight cultures were diluted in fresh media to an OD_600_ of 0.03 and grown at 37 °C with shaking until the mid-log phase. When cultures reached OD_600_ of 0.4-0.5, bacterial cultures were diluted to OD_600_ ∼0.01 in ice-cold PBS and kept on ice until measurement. Flow cytometry was performed using a Beckman Coulter Cytoflex S equipped with a 488 nm excitation laser and a 525/40 nm emission filter. 10,000 events were collected per sample, and 10,000 events of a suspension of calibration beads (Spherotech RCP-30-5A) were collected per flow cytometry run. Analysis of flow cytometry data was performed using the FlowCal software (Castillo-Hair et al., 2016). Each sample of bacteria was density gated on forward and side scatter to retain 70% of events, removing unwanted debris. Similarly, the bead suspension samples were gated to retain 30% of events. The GFP fluorescence of events from bacterial samples was then converted from arbitrary units to Molecules of Equivalent Fluorescein (MEFL) using a standard curve constructed from bead suspension data, as previously described (Gerhardt et al., 2016). The mean of the MEFL values was calculated for each of the three biological replicates.

For plate reader measurements (Figure S2C), overnight cultures were diluted in fresh media to the OD_600_ of 0.03, and were grown in 96 back well with optical bottom (Thermo Scientific) in 200 μl volume at 37 °C with orbital shaking in BioTek Synergy H1 plate reader, and OD_600_ and green fluorescence (excitation, 475 nm; emission, 509 nm) were monitored. The calculation of the GFP expression levels normalized by OD_600_ was done in accordance with (Gao & Stock, 2017) and (Boyer et al., 2010). Specifically, first, the absorbance and fluorescence raw data are fitted by means of cubic smoothing splines, using the csaps function with default parameters of the Spline toolbox in MATLAB. Next, we correct for background fluorescence (Mutalik et al., 2009). The culture background autofluorescence is determined using empty vector strains (AK-1698 and AK-1704 in Table S4). Specifically, the replicate data points of the reporterless strain were averaged and used to generate a standard curve ‘Raw Fluorescence vs OD’. The ‘Raw Fluorescence vs OD’ curve of the reporterless strain was then used to subtract background fluorescence from the reporter strain’s fluorescence value at the same OD. Next, we normalized the background corrected fluorescence values by the corresponding OD values. Finally, for each of the nine replicates we calculated a correlation plot between time and OD values. Based on the correlation we converted the background corrected fluorescence values to be a function of time instead of OD (Figure S2C).

### 4.3 Mathematical model of σ^E^ regulatory network

We developed a mathematical model of σ^E^ regulation to investigate the *E. coli* σ^E^ regulatory network and its role in stress adaptation and recovery. An illustration of the network and model reactions is shown in Figure 1A and Figure 2A, respectively. The model reactions are described in Results Section 2.1 and a summary table containing, model reactions, transcription equations, parameter values used, and their respective references can be found in Table S1. Model ordinary differential equations are shown in Table S2.

### 4.4 Model optimization algorithm

To estimate the unknown model parameters (Table S1), we implemented a model optimization algorithm, incorporating nine constraints to guide the fitting process (Table S3). A description of each constraint is in Results Section 2.1. Constraints C1 and C2 were implemented by directly comparing the steady-state concentrations of RseA and σ^E^ under optimal conditions (*i.e.,* RseA half-life equal to 8 min, (Ades et al., 2003)). Meanwhile, mathematically, given E1 in Table S2, the implementation of C3 is written as:

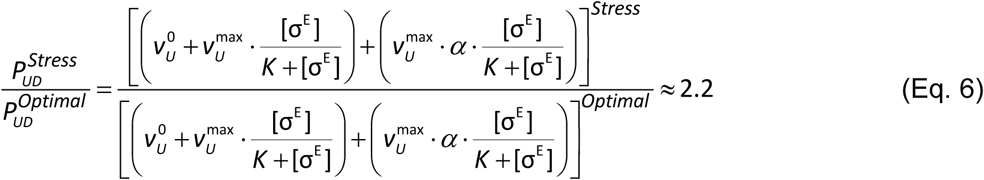

Where, *P_UD_* stands for the sum of the upstream and downstream promoter activities. Similarly, given E2 in Table S2, mathematically, constrain 4 (C4) can be written as:

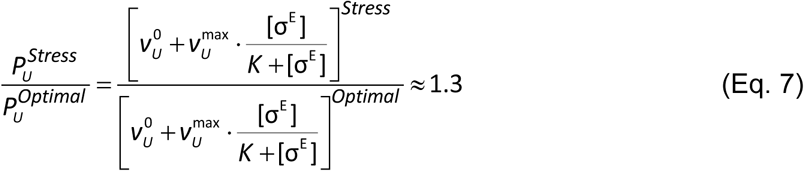

Where, *P_U_* corresponds to the promoter activity of the upstream promoter, the promoter that regulates σ^E^ expression. Constrain 5 (C5) is mathematically implemented in the model by setting:

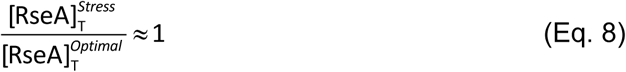

Where, [RseA]_T_ stands for total concentration of RseA. Constrain C6 is implemented by setting the ratio between complex dissociation (k_OFF_) and formation (k_ON_) to be 10 pM (Chaba et al., 2007). Constrain C7, is implemented by comparing the concentration of free σ^E^ under stress conditions at steady-state.

To further constrain the model, we engineered two mutant strains. Specifically, we engineered a strain with a reporter plasmid for the upstream promoter (strain WT, Methods Section 4.1). The second strain, contains a reporter plasmid for the intact upstream promoter. However, the native *rseA* gene was knocked out (Methods Section 4.1). From the plate reader measurements, we calculated the relative promoter activity (Methods Section 4.2) of the strains to be ∼ 3.6. We implement this constraint (C8) in the model by optimizing:

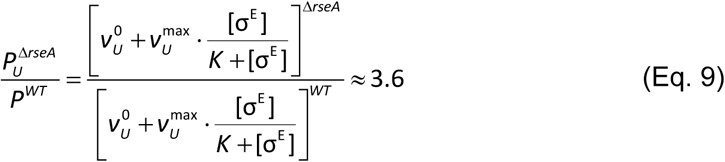

Constrain 9 (C9) is implemented by optimizing the rseA promoter activity (i.e. positive terms in E1 of Table S2) to follow the same dynamics as the RseA synthesis decay shown in Figure S6. RseA synthesis decay was extracted from (Ades et al., 2003) using Graph Grabber 2.0.2 software (Bendow, 2020).

Next, to find the best fitting rates constants that satisfy these constraints, we defined an objective function for optimization. This function minimizes the sum of the squared differences between the constraints and model predictions. Global optimization was performed using the fmincon function in MATLAB R2023b. To minimize the risk of converging to local minima and to assess parameter uncertainty, we additionally implemented an ensemble-based local optimization strategy. This involved randomly sampling 2000 parameter sets within the defined boundaries. The ensemble was screened using MATLAB’s pattern search algorithm and only the 50 parameters sets with the lowest mean squared error were used for optimization and analysis (Ades et al., 2003). While the global optimization yielded a single best-fitting parameter set (Table S1), the ensemble approach enabled us to evaluate parameter uncertainty by analyzing the distribution of well-fitting parameter optimized sets. Overall, the inferred parameter uncertainty was low (Figure S1).

### 4.5 Model simulations

Deterministic model simulations (Figures 2C, 3B-E, 4, 5, 6) were performed using the *ode15s* solver in MATLAB R2023b. In Figures 2C, 3C-E, and 6B–C, we analyzed the transient dynamics of model species prior to reaching steady state. In the other figures, simulations were run for over 1000 minutes to allow the system to reach steady state.

In the model, stress conditions were simulated by dynamically adjusting the overall degradation rate of RseA. Specifically, during the initial stress phase (i.e., the first 10 minutes), the RseA half-life was set to 2 minutes (Ades et al., 2003). In the subsequent adaptation phase, the half-life was increased to 4.5 minutes (Ades et al., 2003). Finally, to simulate recovery from stress, the RseA half-life was increased to 50 minutes (Ades et al., 2003). As estimated by the *model 2.0*, changes in RseA half-life between phases were implemented gradually, except in Figure 2C (yellow line), where it was set to change instantaneously at the onset of the stress response (t = 0 h).

In Figure 3D, the model simulation results were compared to the experimental data from (Ades et al., 2003), extracted using Graph Grabber 2.0.2 software (Bendow, 2020).

## 5. Supplemental data

Supplemental data available online.

## 6. Author contributions

C.S.D.P., A.K., and O.A.I. conceived the study. A.K., and O.A.I. supervised the study. C.S.D.P. executed the model implementation and simulation, to which N.A., and K.L. contributed to. C.S.D.P performed data analysis and results interpretation. M.B., H.D., K.L., and A.K. performed the experimental measurements, except for flow-cytometry measurements and analysis, which were carried out by D.J.H. The manuscript was written by C.S.D.P., N.A., A.K., and O.A.I.

## 7. Acknowledgements

This work was supported by Jenny and Antti Wihuri Foundation [to C.S.D.P.], by National Science Foundation [NSF-PHYS-2019745 to N.A], by the National Science Foundation Graduate Research Fellowship [1842494 to D.J.H], by NIGMS [1R35GM156651 to A.K], and by National Science Foundation [MCB-2204402 to O.A.I (co-PI)]. In addition, we also thank Jeffrey J. Tabor for access to laboratory resources.

## SUPPLEMENTAL MATERIAL

### Supplementary Tables

**Table S1:** Kinetic parameters and model reactions of the mathematical model of σ^E^ regulatory network (Methods Section 4.3). Model rate constants were optimized using the model optimization algorithm described in (Methods section 4.4). The degradation of RseA was assumed to be equal to 0.0015 s^-1^ under optimal conditions (Ades et al., 2003). During stress conditions, the degradation rate of RseA is further reduced (Ades et al., 2003). The mRNA half-life was assumed to be 2.5 min (Bernstein et al., 2002) and the cell doubling time was assumed to take 30 min. The subscripts *U* and *D* in R1 and R2 refer to upstream promoter and downstream promoter, respectively.

| Reaction | Event | Reaction | Rate constants | References |
| --- | --- | --- | --- | --- |
| **R1** | Production of $\sigma^E$ and RseA mRNA<sub>s</sub> from P1 | $v_U \rightarrow e + r$ | $v_U = v_U^0 + v_U^{\max} \cdot \frac{[\sigma^E]}{K + [\sigma^E]}$<br>$v_U^0 = 1.14 \times 10^{-6} \text{ } \mu\text{M} \cdot \text{s}^{-1};$<br>$v_U^{\max} = 8.5 \times 10^{-6} \text{ } \mu\text{M} \cdot \text{s}^{-1};$<br>$K = 0.10 \text{ } \mu\text{M}$ | $v_{P1}^0$, optimized;<br>$v_{P1}^{\max}$, optimized;<br>$K$, optimized; |
| **R2** | Production of $\sigma^E$ and RseA mRNA<sub>s</sub> from P2 | $v_D \rightarrow r$ | $v_D = v_U^{\max} \cdot \alpha \cdot \frac{[\sigma^E]}{K + [\sigma^E]}$<br>$v_U^{\max} = 1.14 \times 10^{-6} \text{ } \mu\text{M} \cdot \text{s}^{-1}; \alpha = 9.66;$<br>$K = 0.10 \text{ } \mu\text{M}$ | $v_{P2}^{\max}$, optimized;<br>$\alpha$, optimized;<br>$K$, optimized; |
| **R3** | RseA mRNA translation | $r \xrightarrow{k_m} r + R$ | $k_m = 0.23 \text{ s}^{-1}$ | optimized |
| **R4** | $\sigma^E$ mRNA translation | $e \xrightarrow{\beta \cdot k_m} e + \sigma^E$ | $\beta = 0.5; k_m = 0.23 \text{ s}^{-1};$ | $\beta$, own data;<br>$k_m$, optimized; |
| **R5** | Complex formation | $\sigma^E + R \xrightleftharpoons[k_{OFF}]{k_{ON}} C$ | $k_{ON} = 0.08 \text{ } \mu\text{M}^{-1} \cdot \text{s}^{-1};$<br>$k_{OFF} = 8.1 \times 10^{-7} \text{ s}^{-1};$ | $k_{ON}$, optimized;<br>$k_{OFF}$, (Chaba et al., 2007); |
| **R6** | Complex dissociation by degradation of RseA | $C \xrightarrow{k_{dR}} \sigma^E$ | $k_{dR} = 0.0015 \text{ s}^{-1};$ | (Ades et al., 2003) |
| **R7** | $\sigma^E$ mRNA degradation | $e \xrightarrow{k_{deg}} \emptyset$ | $k_{deg} = 0.0046 \text{ s}^{-1};$ | (Bernstein et al., 2002) |
| **R8** | RseA mRNA degradation | $r \xrightarrow{k_{deg}} \emptyset$ | $k_{deg} = 0.0046 \text{ s}^{-1};$ | (Bernstein et al., 2002) |
| **R9** | $\sigma^E$ dilution | $\sigma^E \xrightarrow{k_{dil}} \emptyset$ | $k_{dil} = 3.8 \times 10^{-4} \text{ s}^{-1};$ | Assuming doubling time is 30 min. |
| **R10** | Complex dilution | $C \xrightarrow{k_{dil}} \emptyset$ | $k_{dil} = 3.8 \times 10^{-4} \text{ s}^{-1};$ | Assuming doubling time is 30 min. |
| **R11** | RseA degradation and dilution | $R \xrightarrow{k_{dil} + k_{dR}} \emptyset$ | $k_{dil} = 3.8 \times 10^{-4} \text{ s}^{-1};$<br>$k_{dR} = 0.0015 \text{ s}^{-1};$ | Assuming doubling time is 30 min. (Ades et al., 2003) |

**Table S2:** Model ordinary diferential equations (ODEs), assuming the model reactions in Table S1.

| Equation # | ODE |
| --- | --- |
| **E1** | $\frac{d[r]}{dt} = \left( v_U^0 + v_U^{\max} \cdot \frac{[\sigma^E]}{K + [\sigma^E]} \right) + \left( v_U^{\max} \cdot \alpha \cdot \frac{[\sigma^E]}{K + [\sigma^E]} \right) - k_{\text{deg}} \cdot [r]$ |
| **E2** | $\frac{d[e]}{dt} = \left( v_U^0 + v_U^{\max} \cdot \frac{[\sigma^E]}{K + [\sigma^E]} \right) - k_{\text{deg}} \cdot [e]$ |
| **E3** | $\frac{d[\sigma^E]}{dt} = k_m \cdot \beta \cdot [e] + k_{dR} \cdot [C] + k_{\text{OFF}} \cdot [C] - k_{\text{ON}} \cdot [R] \cdot [\sigma^E] - k_{\text{dil}} \cdot [\sigma^E]$ |
| **E4** | $\frac{d[R]}{dt} = k_m \cdot [r] + k_{\text{OFF}} \cdot [C] - k_{\text{ON}} \cdot [R] \cdot [\sigma^E] - k_{dR} \cdot [R] - k_{\text{dil}} \cdot [R]$ |
| **E5** | $\frac{d[C]}{dt} = k_{\text{ON}} \cdot [R] \cdot [\sigma^E] - k_{\text{OFF}} \cdot [C] - k_{dR} \cdot [C] - k_{\text{dil}} \cdot [C]$ |

**Table S3:** Fitting constraints of the mathematical model of the σ^E^ regulatory network (Methods section 4.4). Strain description of constrain C7 can be found in (Methods Section 4.1). The correspondent mathematical equations to establish each constrain in the model are described in (Methods section 4.4). Shown are the model optimization values for both Model 1.0 (i.e., assuming an immediate change in RseA half-life upon stress) and Model 2.0 (i.e., assuming a gradual change in RseA half-life upon stress)

| Constrain | Description | Variable | Reference | Reference value | Model 1.0 optimized value | Model 2.0 optimized value |
| --- | --- | --- | --- | --- | --- | --- |
| **C1** | Basal total [RseA] | $[RseA]_T$ | (Collinet et al., 2000) | 0.33 μM | 0.31 μM | 0.33 μM |
| **C2** | Basal total [$\sigma^E$] | $[\sigma^E]_T$ | (Collinet et al., 2000) | 0.13 μM | 0.13 μM | 0.14 μM |
| **C3** | Fold increase of RseA synthesis due to heat-shock | $\frac{P_{UD}^{Stress}}{P_{UD}^{Optimal}}$ | (Ades et al., 2003) | 2.2 | 1.7 | 1.8 |
| **C4** | Fold increase of RpoE synthesis due to heat-shock | $\frac{P_U^{Stress}}{P_U^{Optimal}}$ | (Raina et al., 1995) | 1.3 | 1.3 | 1.4 |
| **C5** | Relative level of total RseA during stress recovery | $\frac{[RseA]_T^{Stress}}{[RseA]_T^{Optimal}}$ | (Ades et al., 2003) | 1 | 1.53 | 1.4 |
| **C6** | $\sigma^E$ and RseA dissociation constant | $\frac{k_{OFF}}{k_{ON}}$ | (Chaba et al., 2007) | 10 pM | 9.9 pM | 9.9 pM |
| **C7** | Concentration of free $\sigma^E$ under heat-shock conditions | $\frac{[\sigma^E]_T}{4}$ | Set to avoid fits with low levels of free $\sigma^E$ | 0.03 μM | 0.04 μM | 0.03 μM |
| **C8** | $\sigma^E$ and $\sigma^{70}$ promoter activity of $\Delta rseA$ strain relative to the promoter activity of WT strain | $\frac{P_{\sigma^{70};\sigma^E}^{\Delta rseA}}{P^{WT}}$ | Reported here | 3.6 | 3.5 | 3.8 |
| **C9** | Relative RseA synthesis during stress recovery | $\frac{P_{UD}^{Stress}(t)}{P_{UD}^{Optimal}}$ | (Ades et al., 2003) | Blue circles in Fig. 2B | Yellow line in Fig. 2D | Orange line in Fig. 2D |

**Table S4:**
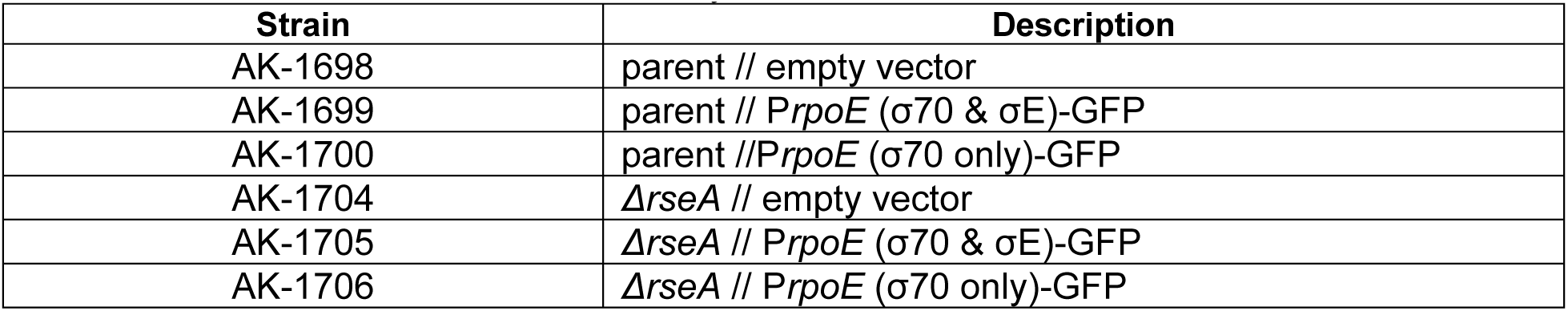
Bacterial strains used in this study.

| Strain | Description |
| --- | --- |
| AK-1698 | parent // empty vector |
| AK-1699 | parent // *PrpoE* (σ70 & σE)-GFP |
| AK-1700 | parent // *PrpoE* (σ70 only)-GFP |
| AK-1704 | ΔrseA // empty vector |
| AK-1705 | ΔrseA // *PrpoE* (σ70 & σE)-GFP |
| AK-1706 | ΔrseA // *PrpoE* (σ70 only)-GFP |

**Table S5:**
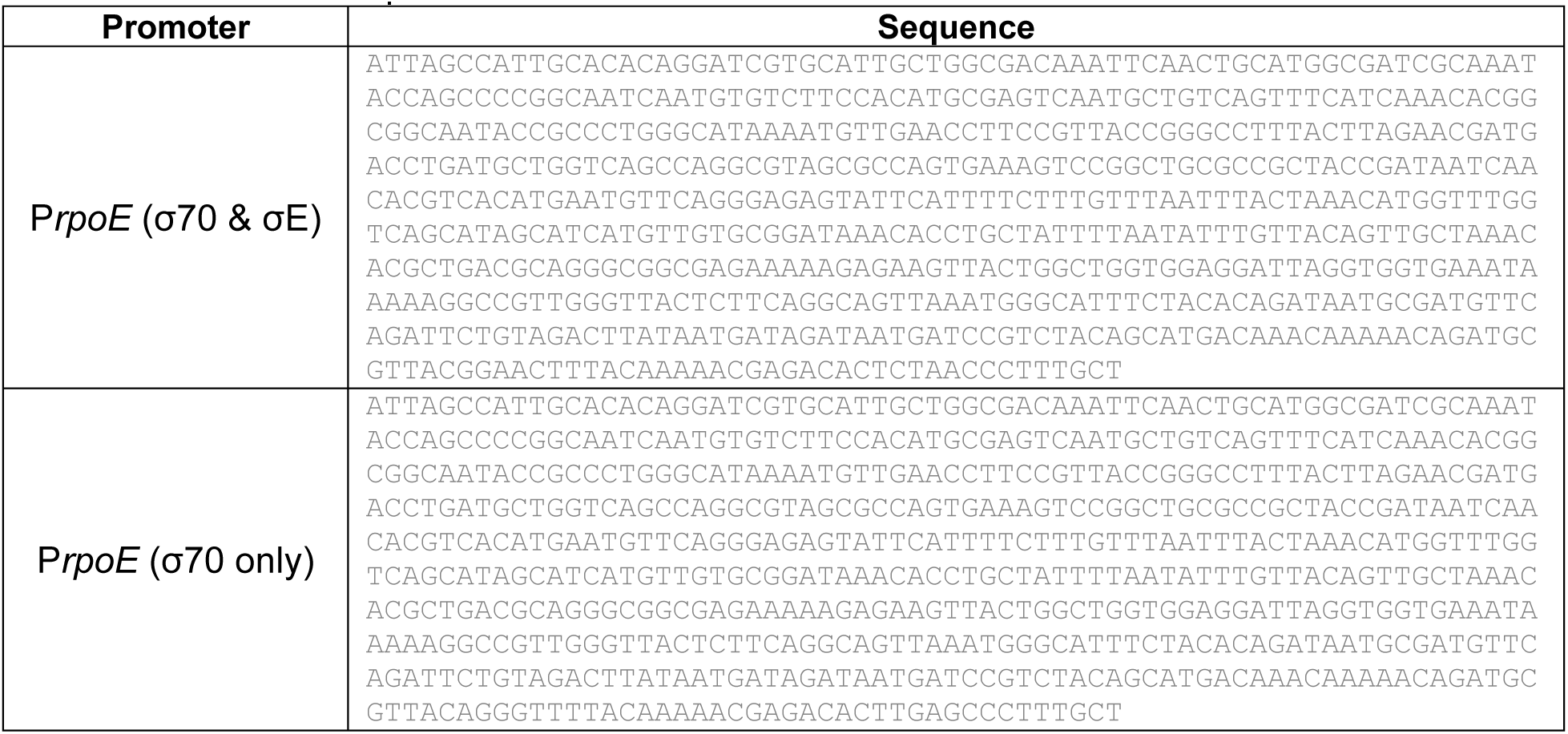
Promoter sequences

### Supplementary Figures

**Figure S1:**
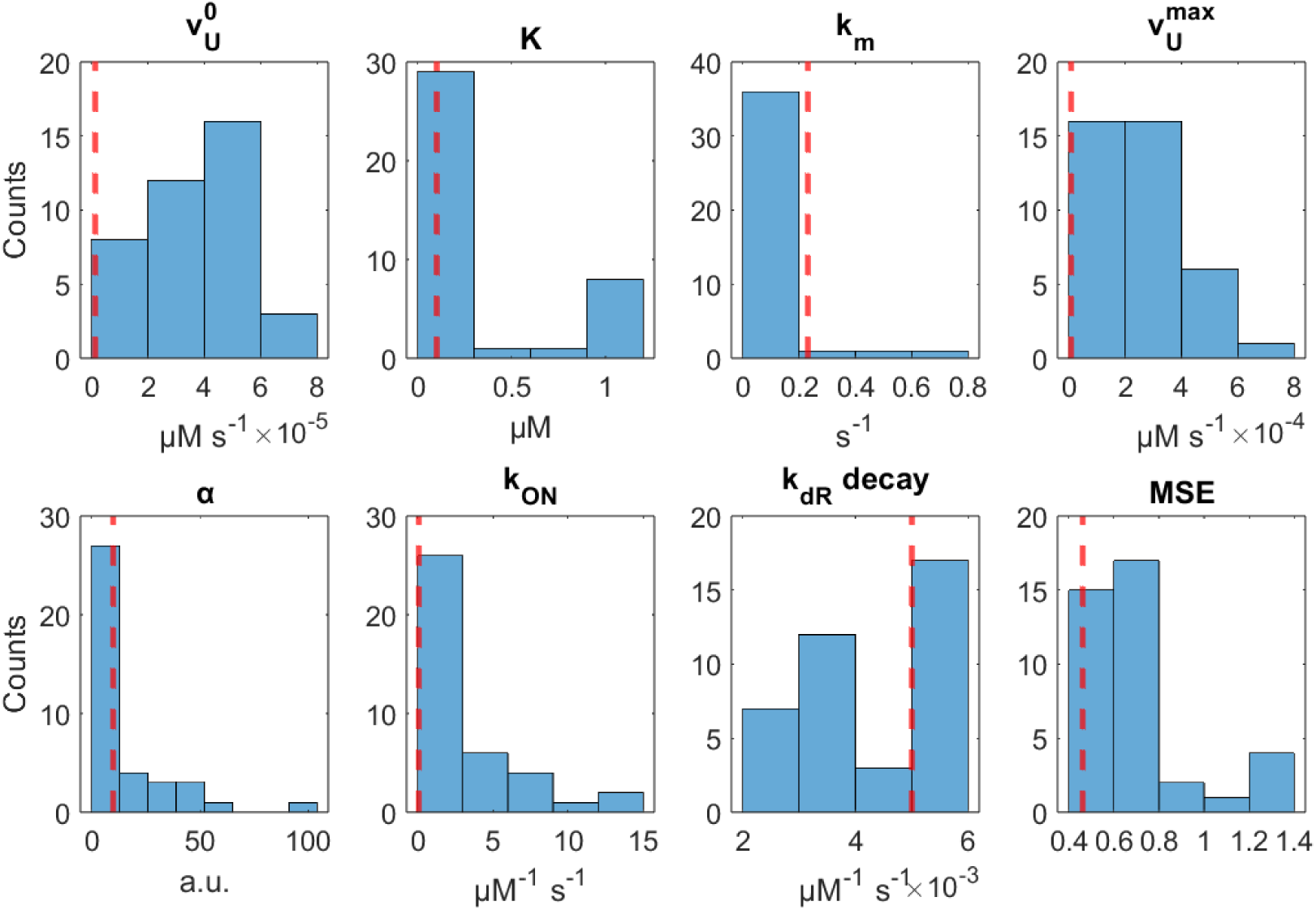
Distribution of fitted parameter values obtained through an ensemble-based local optimization strategy. Shown are the distributions of optimized parameter values from the 50 initial parameters sets with the lowest mean squared error out of the 2000 initially randomly sampled parameter sets (Methods section 4.4). The bottom right graph shows the mean squared error of each parameter set. Vertical dashed red lines correspond to the value of the optimized parameter obtained from the global optimization (Table S1).

**Figure S2:**
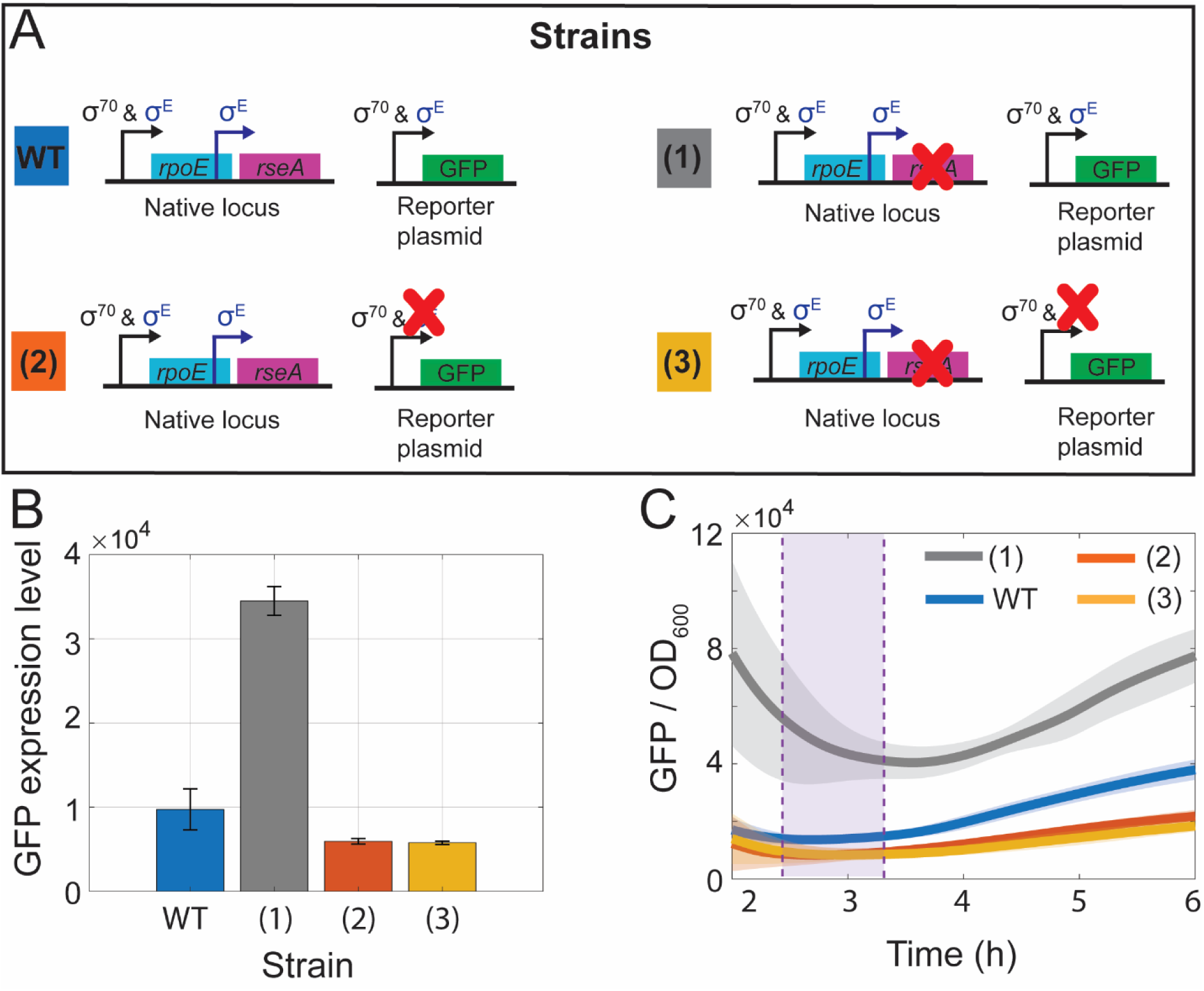
Experimental promoter activity of mutant strains. **(A)** Illustrative representation of the WT strain (WT) and the three other engineered strains. In strain (1) the gene *rseA* was knocked out from the native locus. Strain (2) does not contain promoter specificity to σ^E^ in the reporter plasmid. In strain (3) the gene *rseA* was knocked out from the native locus and the reporter plasmid does not contain promoter specificity to σ^E^. **(B)** GFP expression level measured by flow cytometry, for each strain. Data corresponds to the mean and standard deviation (vertical error bars) of 3 biological replicates. Relative expression of (1)/WT was used to calibrate the model, (Results Section 2.1). Relative expression of (2)/WT and (3)/WT was used to validate model predictions, (Results Section 2.3). **(C)** GFP expression level relative to OD_600_ measured by plate reader. Purple area corresponds to the approximate period in which samples were diluted in PBS for flow cytometry measurements. Data corresponds to the mean and standard deviation (shadow areas) of 9 replicates.

**Figure S3:**
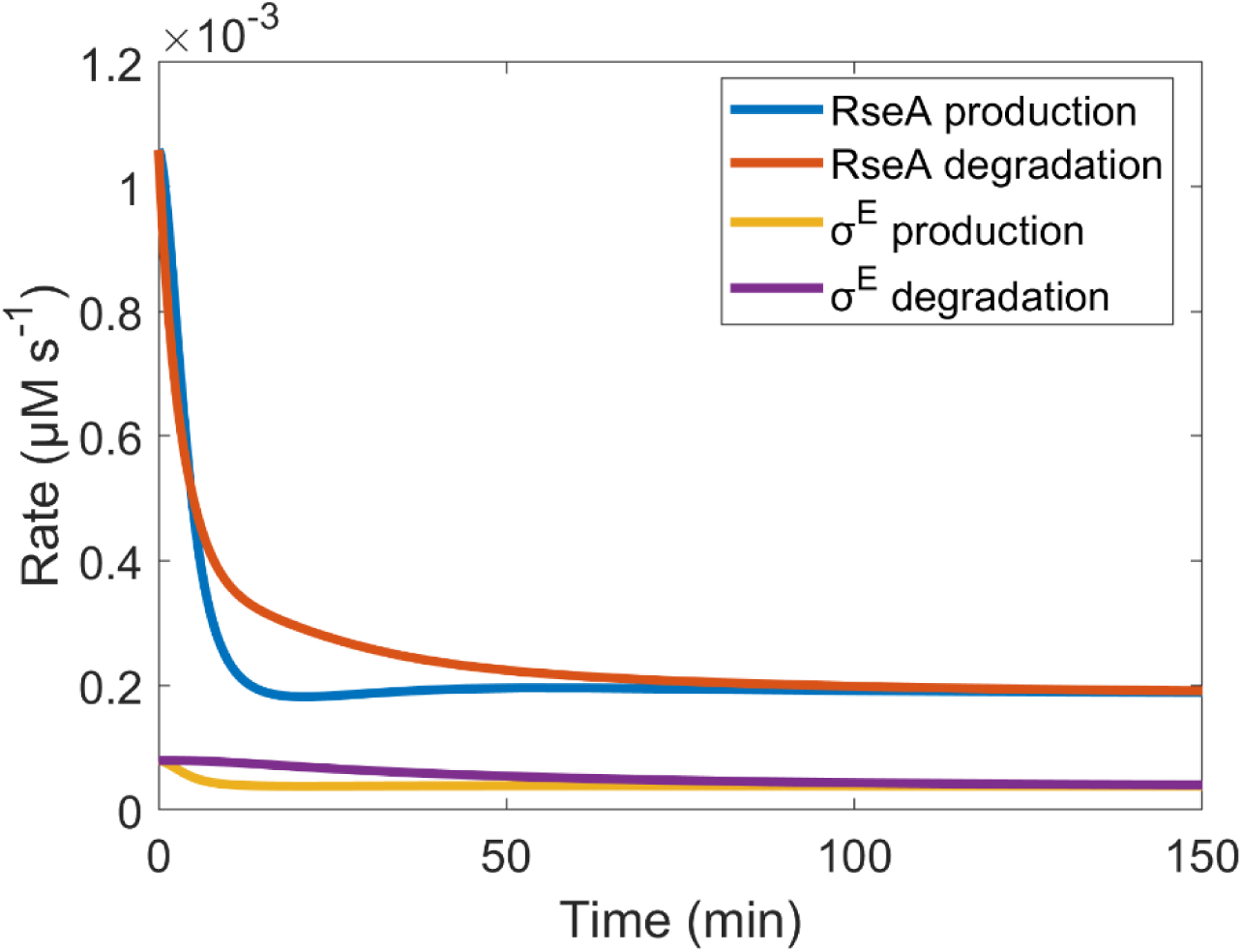
Production and degradation rates of RseA and σ^E^ upon stress recovery. Total RseA production rate was calculated in accordance with 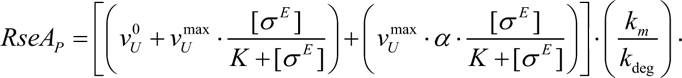 Total RseA degradation rate was calculated in accordance with *RseA_D_* = _(_*k_dil_* + *k_dR_* (*t*))· (*RseA_TOTAL_*). Total σ^E^ production rate was calculated in accordance with 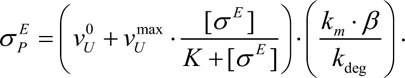 Total σ^E^ degradation rate was calculated in accordance with 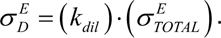

**Figure S4:**
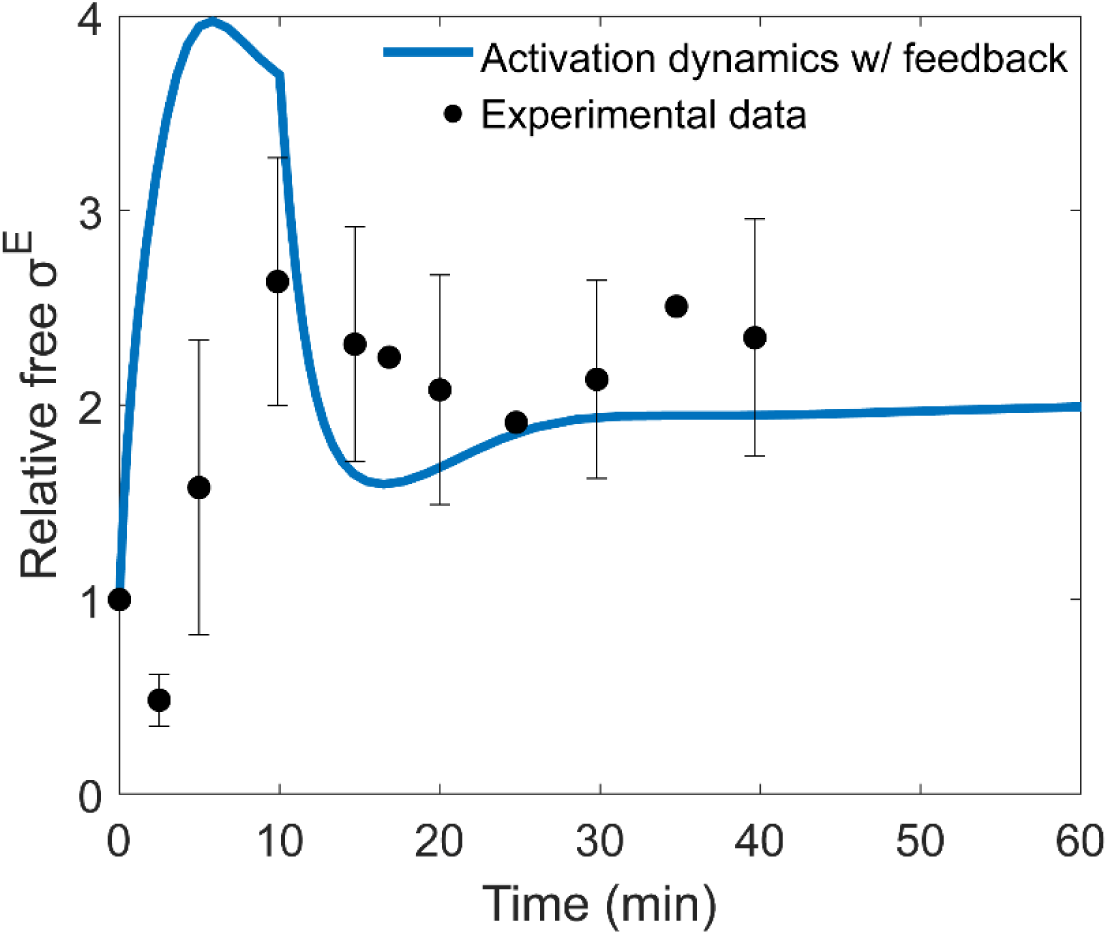
Stress response activation dynamics assuming an immediate decrease in RseA half-life from 8 to 2 minutes, followed by an adaptation phase in which the half-life stabilizes at 4.5 minutes, consistent with observations from (Ades et al., 2003). Experimental data (black dots) are from (Ades et al., 2003).

**Figure S5:**
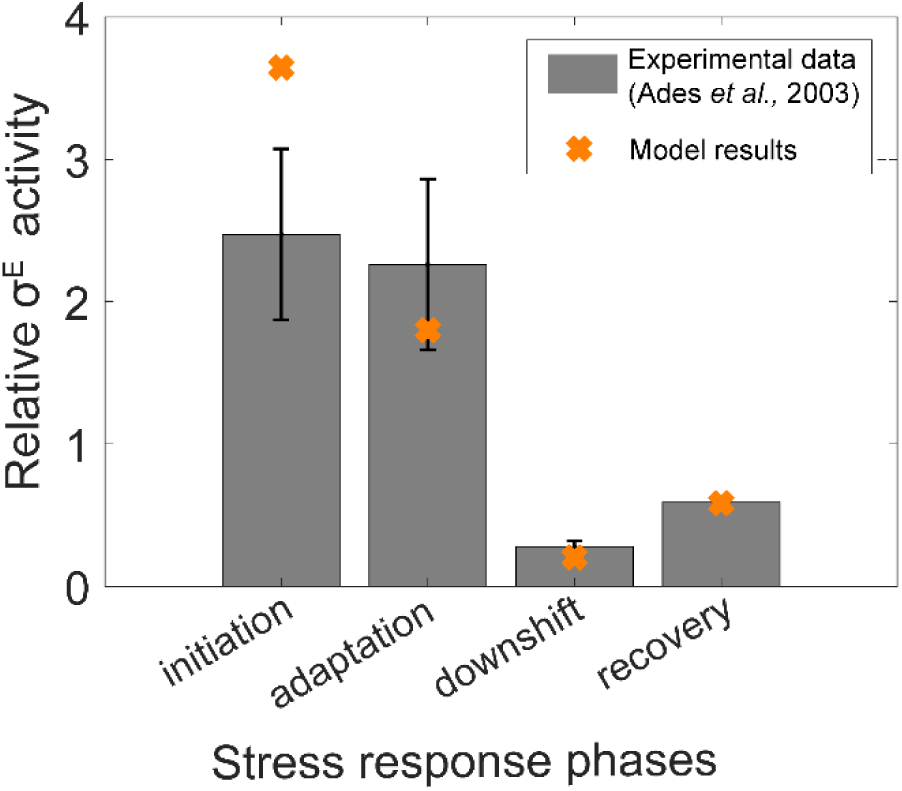
Relative σ^E^ activity levels predicted by the model (orange crosses) compared to experimental measurements from (Ades et al., 2003). We note that the relative σ^E^ activity values for the adaptation and downshift phases of the stress response were used in the model optimization algorithm.

**Figure S6:**
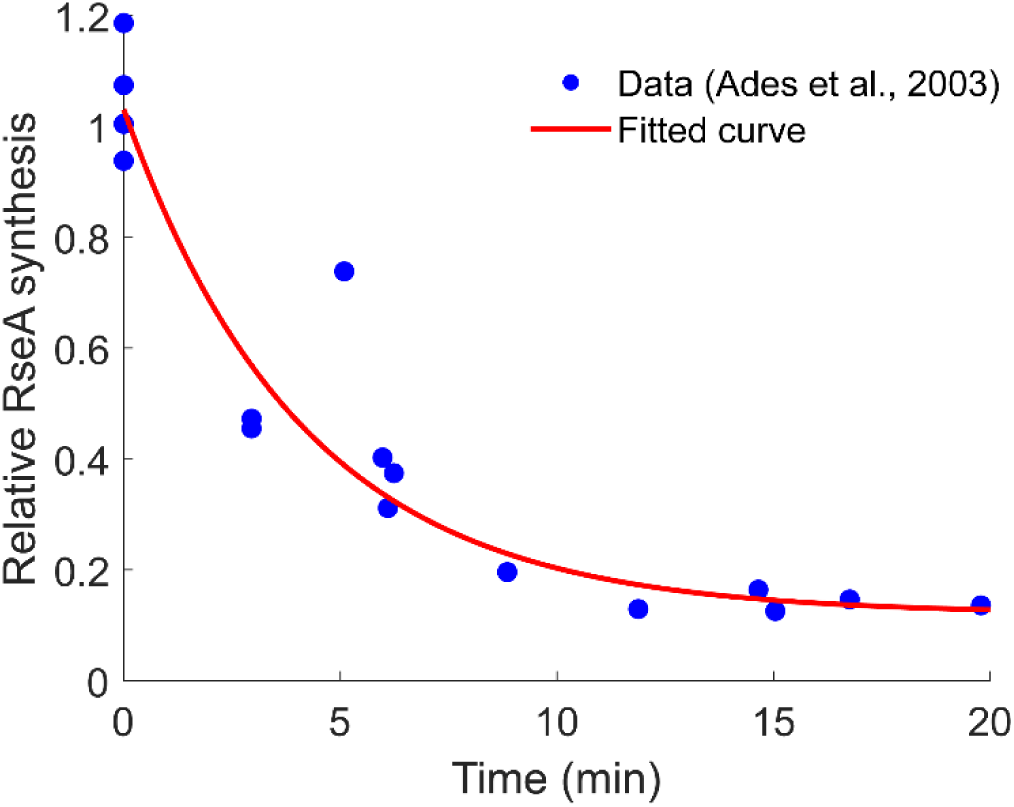
Relative RseA synthesis after temperature downshift from 43°C to 30°C. Blue circles are experimental data reported in (Ades et al., 2003), figure 2A. The red line is the best fit exponential decay (*y*(*t*) = *a*. *e*^−*b*.*t*^ + *c*), estimated from the datapoints. Where *a* = 0.91, *b* = 0.004, and *c* = 0.12.

